# The evolution of the phenylpropanoid pathway entailed pronounced radiations and divergences of enzyme families

**DOI:** 10.1101/2021.05.27.445924

**Authors:** Sophie de Vries, Janine MR Fürst-Jansen, Iker Irisarri, Amra Dhabalia Ashok, Till Ischebeck, Kirstin Feussner, Ilka N Abreu, Maike Petersen, Ivo Feussner, Jan de Vries

## Abstract

Land plants constantly respond to fluctuations in their environment. Part of their response is the production of a diverse repertoire of specialized metabolites. One of the foremost sources for metabolites relevant to environmental responses is the phenylpropanoid pathway, which was long thought to be a land plant-specific adaptation shaped by selective forces in the terrestrial habitat. Recent data have however revealed that streptophyte algae, the algal relatives of land plants, have candidates for the genetic toolkit for phenylpropanoid biosynthesis and produce phenylpropanoid-derived metabolites. Using phylogenetic and sequence analyses, we here show that the enzyme families that orchestrate pivotal steps in phenylpropanoid biosynthesis have independently undergone pronounced radiations and divergence in multiple lineages of major groups of land plants; sister to many of these radiated gene families are streptophyte algal candidates for these enzymes. These radiations suggest a high evolutionary versatility in the enzyme families involved in the phenylpropanoid-derived metabolism across embryophytes. We suggest that this versatility likely translates into functional divergence and may explain the key to one of the defining traits of embryophytes: a rich specialized metabolism.

## INTRODUCTION

A diverse profile of specialized metabolites is one of the characteristics of land plants (embryophytes). Almost any aspect of the biology of land plants is underpinned by specialized metabolites—be it the phytohormones that are major modulators upstream in various regulatory hierarchies (Scheres and van der Putten, 2017; Berens et al., 2017; Blázquez et al., 2020) or pigments that give land plants their color and attune photochemical properties (Jahns and Holzwarth, 2012).

A key aspect of the biological relevance of most specialized metabolites is their use under challenging environmental conditions. Indeed, the elaboration of their specialized metabolism is considered one of the drivers for the massive radiation of embryophytes on land (Weng 2013). Moreover, a diversity of specialized metabolism likely played a key role during the earliest steps of plants on land—allowing for the production of compounds that protected land plants against the challenges of the terrestrial environment such as drought and increased UV radiation (Rensing, 2017; de Vries and Archibald, 2018; Fürst-Jansen et al., 2020; Jiao et al., 2020). A major pathway giving rise to a variety of specialized metabolites that act in warding off environmental stressors is the biosynthesis of phenylpropanoids (Dixon and Paiva, 1995; Dixon et al., 2002; Vogt, 2010).

The phenylpropanoid pathway is the source of precursors for thousands of metabolites with multifaceted functions, and accounts for about 40% of organic carbon on earth (Vogt, 2010). One facet of these functions is that phenylpropanoid-derived compounds act as structural polymers, foremost among which are the different types of lignin (Ralph et al., 2004; Vanholme et al., 2012; Vanholme et al., 2019). Another prominent facet is that these metabolites act as UV-protecting substances. While some of the best-known UV screens are flavonoids, various other compounds stemming from the phenylpropanoid pathway are equally potent UV protectants (Sheahan, 1996; Booij James et al., 2000; Sytar et al. 2018; Xue et al., 2020). The list of links between phenylpropanoid-derived compounds and the response to environmental challenges could be continued; in fact, the response to almost any abiotic stressor that plants face in the terrestrial habitat involves the action of phenylpropanoid-derived compounds (for comprehensive reviews see, e.g., Dixon and Paiva, 1995; Vogt, 2010). Furthermore, phenylpropanoid-derived compounds are involved in the defense responses against plant pathogens in many land plant lineages (Danielsson et al. 2011; Ponce de Leon et al., 2012; König et al., 2014; Overdijk et al., 2016; Carella et al., 2019).

All embryophytes make use of the enzymatic routes in the phenylpropanoid pathway. For example, the utilization of flavonoids under UV stress appears to be a conserved response across Embryophyta (Wolf et al., 2010; Clayton et al., 2018). However, not all embryophytes produce the same compounds under the same stress conditions—in contrary, the diversity of compounds is immense. Major differences in the biosynthesis of phenylpropanoid-derived compounds occur in distinct lineages of land plants. This includes specialized roles such as the flower coloration determining anthocyanins that attract pollinators (Miller et al., 2011; Sheehan et al., 2012); such a role of anthocyanins is obviously limited to flowering plants and can vary even among closely related species (Saito and Harborne, 1992). That said, Piatkowski et al. (2020) phylogenetically inferred that orthologs for the entire anthocyanin biosynthesis pathway are already present in the ancestor of seed plants and more than half of the important orthogroups were already present in the most recent common ancestor of all land plants. An important recent insight into the deep evolutionary roots of flavonoid biosynthesis was the discovery of auronidins—a novel class of red flavonoid pigments that are synthesized in the bryophyte *Marchantia polymorpha* (Berland et al. 2019). Further, for example, Renault et al. (2017a) reported on the enrichment of the *Physcomitrium patens* (moss) cuticle in phenolic compounds—an enrichment that hinges on the action of a cytochrome P450 enzyme that is orthologous to enzymes that act in lignin biosynthesis; the production of lignin might trace its evolutionary roots back to an ancient set of enzymes acting in the production of complex, phenol-enriched polymers (Renault et al., 2019). Carella et al. (2019) showed that the liverwort model plant *Marchantia polymorpha* triggers phenylpropanoid biosynthesis upon attack by the oomycete phytopathogen *Phytophthora palmivora*. Similar responses towards phytopathogens are known from gymnosperms (Oliva et al., 2015) and angiosperms (Dixon and Paiva, 1995; Bednarek et al., 2005; Kaur et al., 2010; Chezem et al., 2017; Carella et al., 2019). Thus, all land plants use the core framework of the phenylpropanoid pathway to produce—often lineage-specific—variations of phenylpropanoid derivatives that aid in response to biotic and abiotic stressors.

The production of the chemical repertoire of land plants is often catalyzed by members of large enzyme-coding gene families (Shockey et al., 2003; Nelson and Werck-Reichhart, 2011; Renault et al., 2017b), and this also seems to be the case for the enzymes involved in phenylpropanoid biosynthesis (Hamberger et al., 2007; Xu et al., 2009; Vogt, 2010). It is thus conceivable that various adaptive forces have shaped the families of enzymes that act in the phenylpropanoid pathway, leading to multiple independent cases of sub-and neofunctionalization (see also Rensing, 2010). An inference of the common (minimal) set of enzymes that were present in the last common ancestors (LCA) of (i) streptophytes, (ii) land plants and their closest streptophyte algal relatives, and (iii) land plants can thus shed light on which enzymatic building blocks evolution acted upon to give rise to the elaborate chassis of the phenylpropanoid pathway.

The phenylpropanoid pathway has long been considered to be specific to Embryophyta. However, homologs of the genes coding for the enzymes that constitute the embryophytic phenylpropanoid pathway can be found in extant algal relatives of land plants, suggesting that they were already present in a common ancestor shared by streptophyte algae and land plants (de Vries et al., 2017; Renault et al., 2019; Maeda and Fernie, 2021). Since the beginning of 2020, we have genome data from all major lineages of Streptophyta—except Coleochaetophyceae (Szövényi et al., 2021); only using this extended repertoire of species and sequences allows us to pinpoint which subfamilies and/or which ancestral enzyme of multiple subfamilies were present in the aforementioned LCAs. Compounds that, in land plants, emerge from the phenylpropanoid pathway are indeed found in algae; these include flavonoids and lignin-like compounds in streptophyte algae (Delwiche et al., 1989; Sørensen et al., 2011; Jiao et al., 2020) and core phenylpropanoid building blocks as well as flavonoids in a phylodiverse set of algae (Goiris et al. 2014). Interestingly, lignin-like compounds were even found in distantly-related red macroalgae (Martone et al., 2009)—although this is likely a case of convergence that builds on an unknown enzymatic framework. However, even within the green lineage (Chloroplastida), the question of the deep evolutionary roots of the phenylpropanoid pathway is wide open.

Investigations of the algal relatives of land plants have strongly benefitted from recent progress in phylogenomics on plants and algae. A major outcome of these recent phylogenomic analyses was that the Zygnematophyceae have been pinpointed as the class of algae most closely related to land plants (Wodniok et al., 2011; Wickett et al., 2014; Leebens-Mack et al., 2019). Hand in hand with these phylogenomic efforts went the generation of genomic (Hori et al., 2014; Nishiyama et al., 2018; Cheng et al., 2019; Wang et al., 2020; Jiao et al., 2020) and transcriptomic data on streptophyte algae (Ju et al., 2015; Rippin et al., 2017; de Vries et al., 2018; de Vries et al., 2020). Additionally, critical gaps in the land plant tree of life have been filled; this includes recent publications of the first genomes of liverworts (Bowman et al., 2017), ferns (Li et al., 2018) and hornworts (Szövenyi et al., 2015; Li et al., 2020; Zhang et al., 2020). These data allow for the fine-grained tracing of the evolution of key plant enzyme families across the green tree of life. Recent studies have illuminated the diversity of enzymes in the routes towards flavonoids and anthocyanins as well as the PAL-dependent pathway of salicylic acid biosynthesis via benzoic acid (Piatkowski et al., 2020; Güngör et al., 2021, de Vries et al., 2021).

In this study, we infer the evolutionary history of eleven critical enzyme families known to be woven into the mesh of routes from phenylpropanoids to lignin biosynthesis in land plants; we have paid particular attention to the routes leading to the biosynthesis of lignin. We use the new diversity of genomic and transcriptomic data from land plants as well as streptophyte and chlorophyte algae to infer the origin of these large gene families. The datasets were chosen in a manner that they cover the breadth of streptophyte diversity while providing a balanced sampling; the latter is especially relevant in light of the high number of genomes of flowering plants. We aimed to include at least one representative of each of the major lineages of streptophytes in the datasets we surveyed. Our data pinpoint deep homologs of candidate enzymes in streptophyte algae for L-**p**henylalanine **a**mmonia-**l**yase (PAL), **4**-**c**oumarate–CoA **l**igase (4CL), **c**affeoyl-**CoA *O***-**m**ethyltransferase (CCoAOMT); further, for streptophyte *and* chlorophyte algae, we pinpoint homologs for **c**innamoyl-**C**oA **r**eductase (CCR), **c**innamyl **a**lcohol **d**ehydrogenase (CAD), and potentially relevant **m**ono**a**cyl**g**lycerol **l**ipases (MAGLs). Further, we find that often the functionally characterized enzymes of the core phenylpropanoid and lignin biosynthesis routes derive from lineage-specific radiations, limiting the inference of function outside the model system. Nonetheless we could infer which subfamilies were present in LCAs along the trajectory of streptophyte evolution, even though ancestral functional inference was limited. That said, for enzyme families with deep homologs, we approximated the function through domain prediction and the conservation (or lack thereof) of key residues of known functional importance. We found that all enzyme families underwent several lineage-specific expansions and losses as well as bursts in growth of enzyme families that occurred early during the radiation of land plants. We hypothesize that lineage-specific expansions in these enzyme families is linked with the diversity of lineage-specific phenylpropanoid derivatives and functions that occur in the species analyzed here.

## RESULTS AND DISCUSSION

### The checkered occurrence of phenylalanine ammonia-lyase among streptophyte algae

The conversion of the aromatic amino acid phenylalanine and/or tyrosine into cinnamate and/or *p*-coumarate is the first step of the plant phenylpropanoid pathway (Figure 1). This first committed step is catalyzed by PAL and the bifunctional L-phenylalanine/ L-tyrosine ammonia-lyase (PTAL; Barros and Dixon, 2020). For a long time, it was thought that among Chloroplastida, PAL/PTAL were limited to land plants; their gain was considered to have occurred via a lateral gene transfer event that has occurred at the base of the land plant clade (Emiliani et al., 2009). Recently, however, genes coding for putative PAL-like enzymes were detected in streptophyte algae, such as the filamentous streptophyte alga *Klebsormidium nitens* (de Vries et al., 2017). In light of the recent surge in available genomes from across the green tree of life, we set out to explore the evolutionary history of PAL.

**Figure 1.**
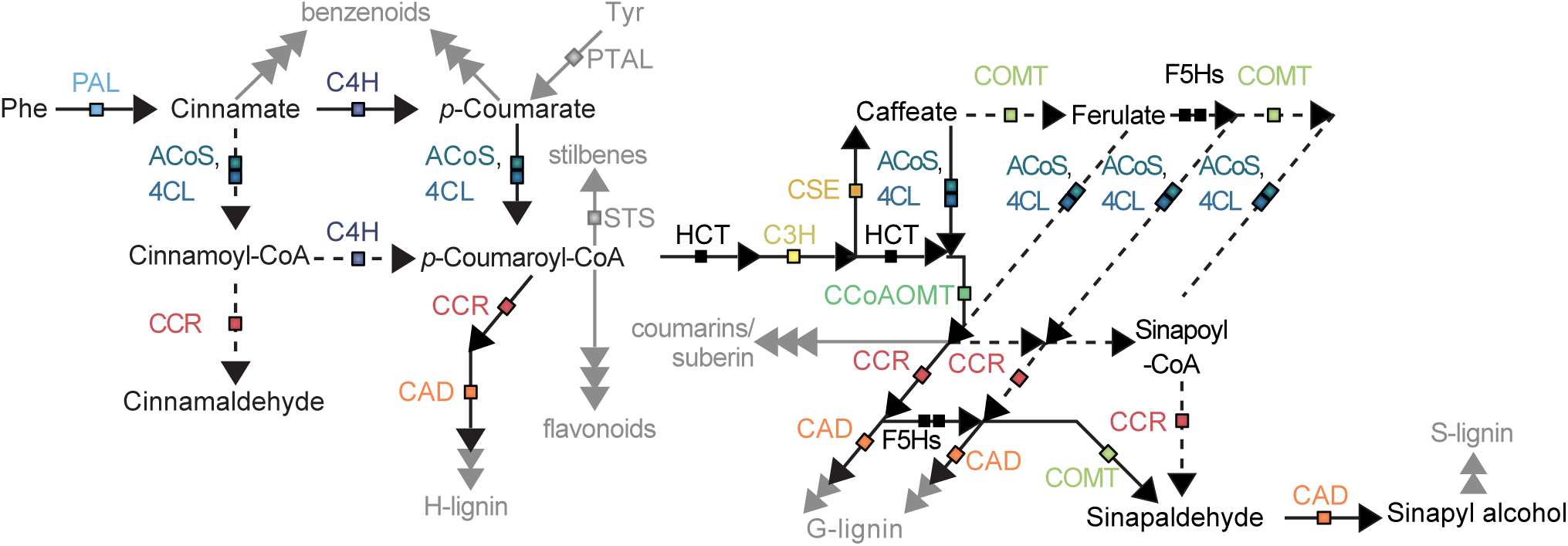
Enzymes involved in the biosynthesis of phenylpropanoid-derived compounds investigated here. A simplified schematic of the phenylpropanoid pathway and its routes to different derivatives are shown. Boxes indicate enzyme families, which are mentioned above each box and color-coded. Their coloration is the same as in Figure 10. Dotted lines indicate putative steps in the pathway.

Using *At*PAL1 as a bait sequence, we screened protein data from diverse land plants and all streptophyte algal genomes available. Hits among streptophyte algae fell into three categories: (i) proteins of between 480 (*Klebormidium nitens* PAL, kfl00104_0290_v1.1) and 527 amino acids (two homologs in *Chara braunii*; g57646_t1 and g34530_t1); (ii) short proteins such as ME1156409C09523 of *Mesotaenium endlicherianum*, which is 184 amino acid in length; (iii) long proteins of between 991 (*Chlorykbus atmophyticus* Chrsp482S06115) and 1115 amino acids (*Klebsormidium nitens* kfl00024_0250_v1.1). Proteins falling into the third category are fusions of an aromatic amino acid lyase domain and a putative tRNA synthetase; homologs of such also occur in land plants (e.g., AT3G02760 and Os05g0150900). The short proteins of category ‘(ii)’ are found in some Zygnematophyceae and are proteins of unknown function with a putative HAL domain. Genomic artefacts leading to this result can be excluded given that these types of protein-encoding genes have been recovered for independent Zygnematophyceae that are likely >500 million years divergent from one another. The proteins in category ‘(i)’ are those with the highest identity to *bona fide* land plant PALs. This category includes the promising PAL candidate kfl00104_0290_v1.1 (see de Vries et al., 2017). We therefore set out to further explore these PAL-like candidates, which noteworthily were only found in the genome of *Klebsormidium nitens* and *Chara braunii* and no other streptophyte algal genome data.

To understand the evolutionary history of PAL in streptophytes, we computed a maximum likelihood phylogeny (Figure 2). The phylogenetic analysis included the aforementioned PAL homologs from diverse Streptophyta as well as bacterial and fungal PALs. In agreement with previous studies (Emiliani et al., 2009; de Vries et al., 2017), the fungal and bacterial PAL sequences are closely related (bacterial clade: bootstrap support 100; fungal clade: bootstrap support 85) to the clade of land plant PALs (bootstrap support: 96). Additionally, we included diverse eukaryotic and prokaryotic histidine ammonia lyases (HALs) based on the set obtained from de Vries et al. (2017), which, as in this latter study, form two clades with eukaryotic and prokaryotic HALs (bootstrap support 83 and 97). All putative PAL-like candidate sequences from *Chara braunii* clustered with HAL sequences, one (*Chara braunii* g66119_t1) with a cyanobacterium (bootstrap support 100), and two (*Chara braunii* g34530_t1 and *Chara braunii* g57646_t1) sister to an entire bacterial HAL clade (bootstrap support 100). Both sequences retrieved for *Klebsormidium nitens*, in contrast, clustered with the PAL clades, one *K. nitens* kfl00024_0250_v1.1 with low support (boostrap level 61) and a rather long branch as sister to plant and fungal PALs, showing that its placement is not fully resolved and that further analyses are required to identify its true identity. The second *Klebsormidium nitens* sequence, kfl00104_0290_v1.1, clustered within the clade of bacterial PALs (bootstrap support 100), of which some were already functionally characterized—for example, the characterized PAL of *Nostoc punctiforme* (Moffit et al., 2007). This is in agreement with the placement of this protein sequence in de Vries et al. 2017 and supports it as a putative PAL sequence.

**Figure 2.**
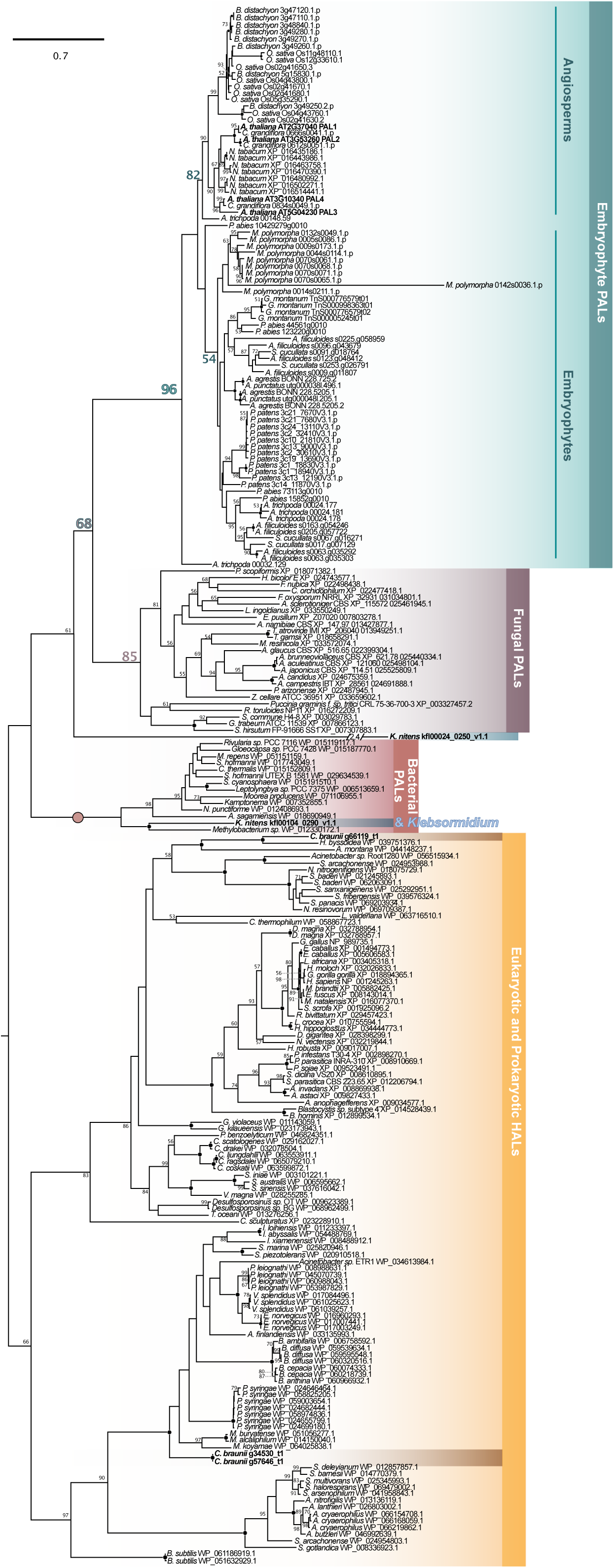
A phylogenetic framework for the origin of streptophyte PAL. PAL and HAL homologs were screened for in fifteen land plant, seven streptophyte algae, and five chlorophytes. Among Chloroplastida, PAL homologs were only recovered from genomes of land plants and the streptophyte algae *Klebsormidium nitens* and *Chara braunii*. From all detected homologs, a rooted maximum likelihood phylogeny was computed using LG+I+G4 as model for protein evolution (chosen according to BIC). 100 bootstrap replicates were computed; only bootstrap values ≥50 are shown and bootstrap values of 100 are depicted by a filled dot. Colored font and dots correspond to the support recovered for the higher-order clades labeled on the right of the phylogenies.

In sum, the evolutionary origin of streptophyte PAL appears to be complex and remains obscure: it may be that PAL had a distinct origin in streptophyte algae and land plants, yet the pattern may also be explained by an origin via an endosymbiotic gene transfer from the cyanobacterial plastid progenitor that was retained by streptophytes with a gain of an extra C-terminal domain later in the evolution of embryophytes, resulting in the two distinct PAL clades. It is however important to note that the 3’ region of the genomic locus that codes for the shorter *K. nitens* protein kfl00104_0290_v1.1 contains sequence information that resembles code for the missing C-terminal stretch; thus, the C-terminal stretch might have simply been secondarily lost in *K. nitens*. Independently, the presence of PALs in fungi further complicates the evolutionary scenario. Rampant gene losses during eukaryotic evolution or convergent domain acquisitions, as well as horizontal gene transfer from plants (as hypothesized by Emiliani et al., 2009) are other scenarios that can explain the evolutionary origin of PALs in streptophytes and thus ultimately in land plants.

### Streptophyte algae have an expanded and divergent repertoire of cytochrome P450 monooxygenases with no clear C4H orthologs

After the synthesis of cinnamate by PAL, two routes open up (Figure 1). One of them is the conversion of cinnamate into *p*-coumarate, which is catalyzed by **c**innamate **4**-**h**ydroxylase (C4H). C4H belongs to the large class of CYP450 enzymes, which are present among all domains of life (Omura, 1999). Among the CYP450 enzymes C4H belongs to the CYP450 subfamily 73 (CYP73). In land plants, CYP450s have undergone massive duplication and subfunctionalization, underpinning the specialized metabolic capabilities of embryophytes (Nelson and Werck-Reichhart, 2011); for example, the CYP73 subfamily belongs to the larger CYP71 clan. The specific CYP450 monoxygenases that fall into the group of C4H appear to be limited to land plants: clear orthologs can be found in bryophytes and tracheophytes (Emiliani et al., 2009; de Vries et al., 2017). That said, the product of the reaction carried out by C4H in land plants (*p*-coumarate) has been detected via UHPLC-MS/MS in phylodiverse algae (Goiris et al., 2014). Therefore, there appears to exist a route towards *p*-coumarate that is either independent of C4H via direct transformation of tyrosine by PTAL or carried out by a highly divergent C4H homolog. PTALs have so far however been observed in monocots (Barros et al., 2016; Barros and Dixon, 2020), suggesting a different CYP73 subfamily enzyme that may carry out the reaction. Owing to the recent increase in genomic data available for streptophyte algae, we revisited the question of when C4H-based *p*-coumarate might have emerged and explored CYP450 evolution.

We sampled C4H homologs from seven land plant genomes that had a BLAST bit score (a normalized alignment score) of at least 200, as well as seven streptophyte algal and five chlorophyte algal genomes that had a bit score of at least 100. We aligned all C4H homologs and computed a maximum likelihood phylogeny (Figure S1). The well-characterized C4H of *Arabidopsis* fell into a clade with full (100 out of 100) bootstrap support; this clade included at least one C4H homolog from each of the other six land plants genomes, corroborating the notion that all land plants have C4H orthologs, which appear conserved in their function from bryophytes to tracheophytes (Russel and Conn, 1967; Urban et al., 1994; Ro et al., 2001; Wohl and Petersen, 2020) and thus since the LCA of land plants. However, no algal sequences fell into this clade. That said, we observed four well-supported clades of streptophyte algal CYP450 enzymes (Figure S1). Investigating the genetic distances, we find that some of the streptophyte algal sequences have a closer genetic distance to the C4H-like clade than to sequences from land plants (including *Arabidopsis thaliana*) from other CYP450 subfamilies (Table S1). While these sequences remain of unknown function, they are candidates for the CYP450 enzyme family that catalyzes the C4H-function in algae.

### A deep split of streptophyte 4CL/ACS

The second route that opens up after the PAL-dependent step is the conversion of cinnamate into cinnamoyl-CoA. This is carried out by the AMP-forming synthetase/ligase 4CL and potentially other enzymes annotated as acyl-CoA synthetases (ACS/ACoS) (Shockey et al., 2003; Figure 1). Altogether, these enzymes belong to a large family of distantly related acyl-activating enzymes (AAEs), such as the long-chain acyl-CoA synthetases (LACS) and many more (Shockey et al., 2003; Figure S2). Homologs with affinity to 4CL appear to occur across chlorophytes and streptophytes (Labeeuw et al., 2015). At least in *Arabidopsis*, the family of 4CLs has expanded and includes four canonical (“4CL”) and nine additional 4CL-like (“4CLL”) members, falling into AAE clade IV and V as defined by Shockey et al. (2003). We thus set out to understand what the 4CL repertoire of the last common ancestor of land plants and the one shared with algae might have looked like.

In order to trace the radiation of 4CLs across the green tree of life, we sampled 4CL homologs from genomes of nine land plants, seven streptophyte algae (plus four transcriptomes of streptophyte algae) and five chlorophyte algae that had a minimum of 400 and a maximum of 1150 amino acids in length and showed affinity to the 4CL clade in a larger phylogenetic survey (Figure S2). We recovered a large clade (bootstrap support 85) that included all *bona fide* 4CL paralogs and ACOS5 of *Arabidopsis thaliana* (Figure 3); ACOS5 has been previously associated with the *bona fide* 4CL clade (Shockey et al., 2003) but it did show only inconsistent activity on typical substrates of 4CL (Costa et al., 2005) and appears to have a very specific function in sporopollenin biosynthesis of pollen (de Azevedo Souza et al., 2009). These observations agree with the presence of different (but conserved) amino acids at sites that bind hydroxycinnamate in typical 4CLs (Figure 3), which might suggest a different natural substrate for ACOS given that affinity is mostly determined by the binding pocket size (Hu et al. 2010). We further recovered the angiosperm-specific separation defined by Ehlting et al. (1999) into class I and class II 4CLs. Additional lineage-specific radiations occurred, for example in *Physcomitrium patens*, which fell into a clade of bryophyte sequences (bootstrap support of 70) and in *Selaginella moellendorffii* (spread out over the fully-supported clade of 4CLs). The common ancestor of land plants appears to have possessed an ACOS5-like and one 4CL-like gene, all other 4CL paralogs in this clade likely emerged later during land plant evolution. Clustering with AAE clade IV (including *At*ACOS5, and *At*4CL1,2,3 and 5 (bootstrap support 85)) are sequences from five streptophyte algae. Each of the five streptophyte algae possesses one homolog to these five types of AMP-forming ligases with 4-coumarate–CoA synthesizing activity. When we predicted the tertiary structure of *Chlorokybus atmophyticus* Chrsp175S02417 and *Penium margaritaceum* 006213.t1 via Iterative Threading ASSEmbly Refinement (I-TASSER; Zhang 2008), we recovered, in both cases, firefly luciferases as best match (TM-scores 0.852 and 0.918; 1BA3 and 2D1S; Franks et al., 1998; Nakatsu et al., 2006). Investigating the putative structure of the other streptophyte algal sequences (*Zygnema circumcarinatum* DN42558_c0_g1_i1, *Spirogyra pratensis* 3442_c2_g1_i6, and *Klebsormidium nitens* 00016_0470_v1.1), however, always recovered *Populus tomentosa* 4CL (3A9U; Hu et al., 2010) as their closest structural analog (TM scores of 0.957, 0.969, and 0.961, respectively). Hence, we hypothesize that a 4CL/ACOS5-like encoding gene was present in the last common ancestor of all streptophytes. Underpinning this hypothesis is that the sequence of the amino acids in the binding pocket in the streptophyte algal 4CL homologs is consistent with that of 4CL homologs from other land plants (including that of *Arabidopsis thaliana*; Figure 3). Further, the amino acids relevant for the enzymatic function (i.e., the residues KQK involved in adenylation, nucleophilic substitution and coumaroyl-AMP cleavage) are also conserved across most 4CL/ACOS5 sequences, including those of the streptophyte algae. Variation in these residues is already apparent in 4CLL homologs and outside of the 4CL/ACOS5/4CLL clade these residues show high variability (Figure 3).

**Figure 3.**
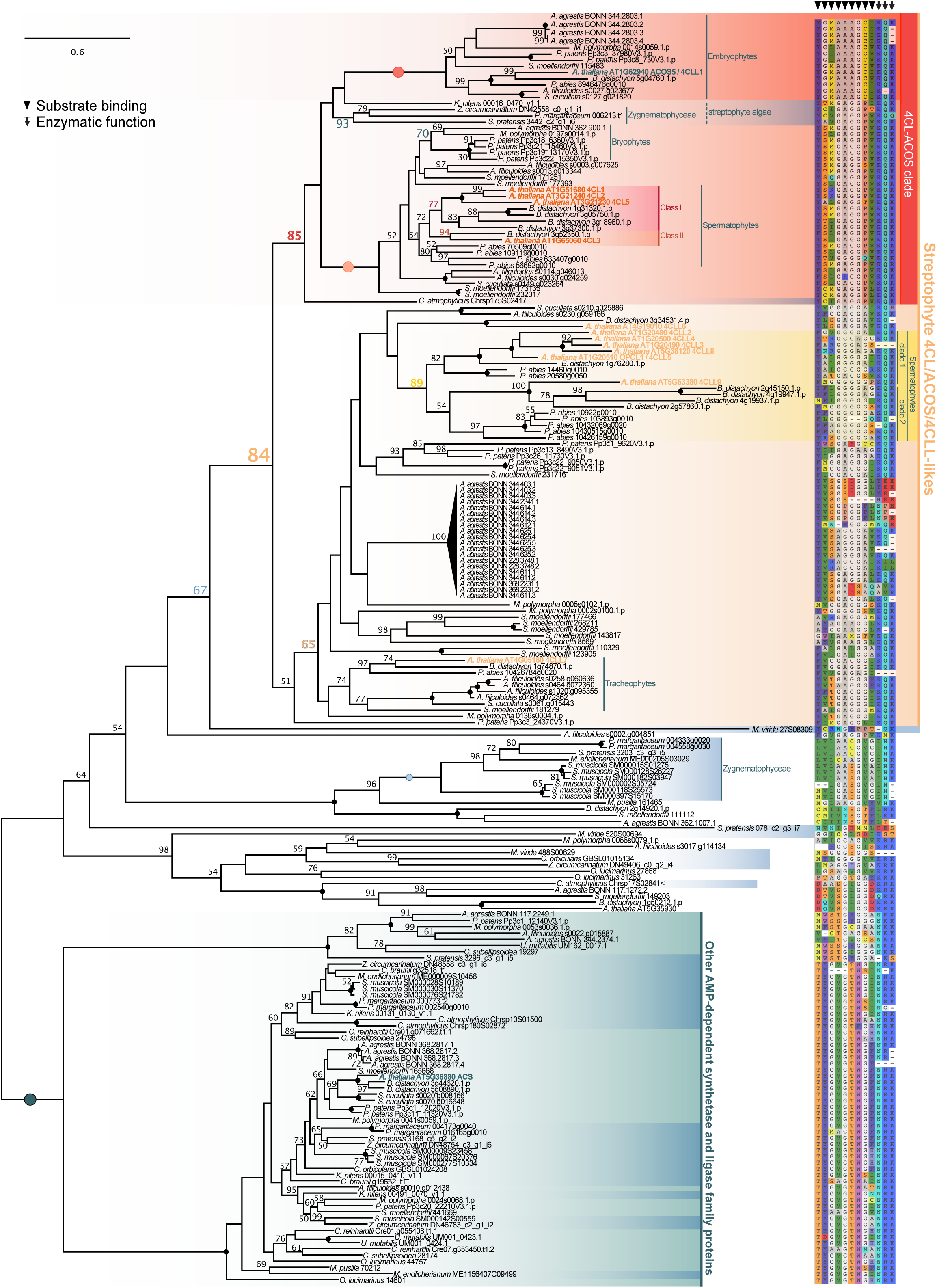
4CL homologs occur across Streptophyta. 4CL homologs were sampled from protein data of nine land plant, seven streptophyte algal and five chlorophyte algal genomes. Only protein sequences with a minimum length of 400 and a maximum length of 1150 amino acids were included. From all detected homologs, a rooted maximum likelihood phylogeny was computed using LG+G4 as model for protein evolution (chosen according to BIC). 100 bootstrap replicates were computed; only bootstrap values ≥50 are shown and bootstrap values of 100 are depicted by a filled dot. Colored font and dots correspond to the support recovered for the higher-order clades labeled on the right of the phylogenies. On the right we show key residues for substrate binding and function of canonical 4CL as reported by Hu et al. (2010).

A similar pattern was observed when we investigated the domain structure of all recovered sequences. Most of the 4CL/ACOS5-like sequences contained four domains: Phosphopantetheine binding ACP domain (IPR025110), AMP-binding, conserved site (IPR020845), AMP-dependent synthetase-like superfamily (IPR042099), AMP-dependent synthetase/ligase (IPR000873; Figure S3). There were four exceptions to this pattern. They include one sequence from the water fern *Azolla filiculoides* (Azfis0013.g013344) and two hornwort sequences from *Anthoceros agrestis* BONN (Sc2ySwM344.2803.3 and Sc2ySwM344.2803.4), which all missed the Phosphopantetheine binding ACP domain (IPR025110). The other exception was *At*ACOS5, which is the only sequence in this clade that missed the conserved AMP-binding site (IPR020845). The domain pattern is similar across the 4CLL-like clade, too. Yet, more sequences miss either the IPR025110 and/or the IPR020845 domain. The streptophyte algal sequences within the ACOS5/4CL clade contained all four domains, whereas algal sequences outside of this clade missed at least one—but recovered several other domains. These additional domains are not conserved within the phylogenetic subclades of these algal sequences and only exceptionally occur in the *4CL/ACOS4/4CLL* (two sequences) or *Other AMP-dependent synthetase and ligase family protein* clades (two sequences).

We recovered a second clade of spermatophyte sequences (bootstrap support 89) representing AAE clade V enzymes, which contains 4CLL8 and several other ATP-ligases of *Arabidopsis thaliana* with predicted 4CL activity, including OPCL1 (matching 4CL-like 5 with a 100% amino acid identity according to Uniprot). Sequences in this clade, however, diverge in the amino acids that are involved in the formation of the binding pocket in the canonical 4CLs (Figure 3), which might point to a different substrate preference of the enzymes in this clade. In fact, OPCL1 and many of these “4CLLs” (e.g., AT5G63380, AT1G20500, AT4G05160) showed higher activity on fatty acids and fatty acid-derived precursors for the phytohormone jasmonic acid than cinnamate-derived compounds in an *in vitro* substrate survey carried out by Kienow et al. (2008). It is thus questionable that the enzymes of this clade act as *bona fide* 4CLs. Homologs to these sequences are found in *Brachypodium distachyon* and *Picea abies,* suggesting an origin in the last common ancestor of seed plants followed by two duplication events with either (a) both taking place in the last common ancestor of angiosperms or (b) one early on in the LCA of seed plants and the second in the LCA of angiosperms. Bootstrap support to include the *Picea abies* sequences in the clade containing AT5G63380 (4CLL9) is however low (bootstrap 54). Each duplication event was followed by independent lineage-specific radiations giving rise to a whole plethora of possible candidates for 4CL, but also a large evolutionary potential with regard to substrate specificity and flexibility. The 4CLL clade of spermatophytes is nested within a larger, lowly supported clade (bootstrap 65) that included sequences from across the land plant tree of life. Here, pronounced and independent expansion occurred in most of the major lineages of land plants, leading to large clades of, for example, proteins of the hornwort *Anthoceros* and the lycophyte *Selaginella*. As noted above, most of the *Selaginella* and some of the *Anthoceros* homologs retained the conserved KQK residues required for the catalytic activity but others did not (Figure 3), which suggests the presence of species-specific functions. Outside of the entire 4CL-ACOS and 4CLL clade (‘Streptophyte 4CL/ACOS/4CLL-likes’; bootstrap 84) clustered various highly divergent ATP-dependent synthetases and ligases that exist throughout the green tree of life including sequences from chlorophytes. None of these synthetases and ligases retained the catalytic triad KQK.

Altogether, our phylogenetic data indicated that a 4CL/ACOS5-like encoding gene was present at the base of Streptophyta. Domain annotation and the analysis of amino acid patterns in the binding pocket and functional sites support this idea. Further, the similarity of these residues between the candidates of streptophyte algal homologs for 4CLs and the sequences of 4CL proteins with high activity on cinnamate derivatives as substrates (see also Costa et al., 2005), indicates that 4CL activity may evolved more than 700 million years ago in streptophytes.

### Patchy distribution of CCR-likes in streptophyte algae and massive independent radiations in land plants

*En route* to the production of different lignin monomers is the NADPH-dependent reduction of the activated acyl-group of the phenylpropanoid backbone molecules. This first step towards an aldehyde functionality is carried out by CCR (Figure 1), which falls into a larger family of NADPH-dependent reductases, including dihydroflavonol reductases (DFRs) and DFR-likes (DFL) (Lacombe et al., 1997; Devic et al., 1999). We previously reported the presence of CCR-like protein sequences in streptophyte algae (de Vries et al. 2017). Since these previous analyses, however, genome data on additional major lineages of land plants and streptophyte algae have become available.

With these new data at hand, covering most major lineages of streptophyte algae and all major lineages of land plants, it is now possible to infer the evolutionary history of CCR-like and DFR-like sequences. We computed a maximum likelihood phylogeny of CCR homologs with a minimum of 220 amino acids that we detected in genomes of 15 land plants, seven streptophyte algae, and five chlorophytes; additionally, we included sequences found in the transcriptomes of the Zygnematophyceae *Spirogyra pratensis* (de Vries et al., 2020), *Zygnema circumcarinatum* (de Vries et al., 2018), and the Coleochaetophyceae *Coleochaete orbicularis* (Ju et al., 2015; Figure 4).

**Figure 4.**
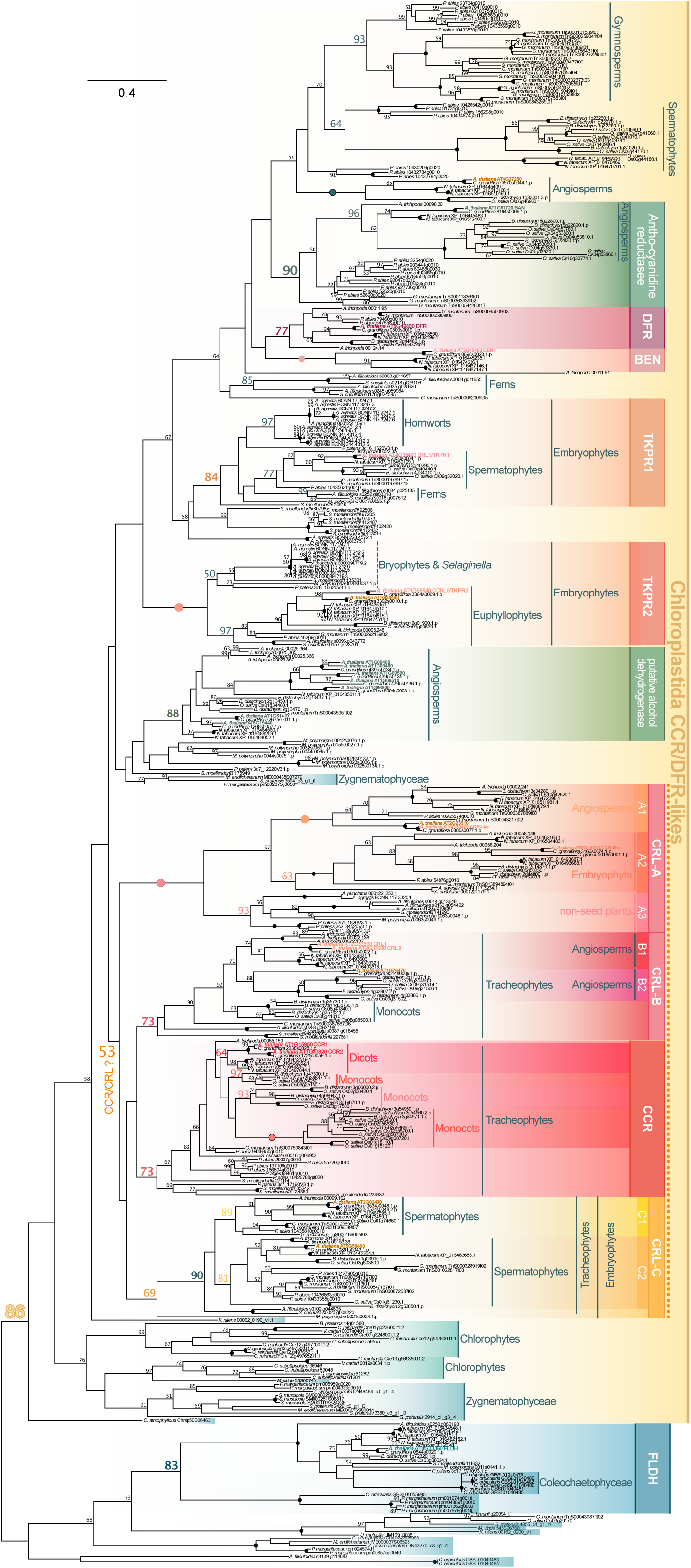
The complex evolutionary history of CCR in Chloroplastida. CCR homologs of a minimum of 220 amino acids were sampled from protein data of 15 land plant, seven streptophyte algal, and five chlorophyte algal genomes; additionally, we included sequences found in the transcriptomes of *Spirogyra pratensis* (de Vries et al., 2020), *Zygnema circumcarinatum* (de Vries et al., 2018), and *Coleochaete orbicularis* (Ju et al., 2015). From all detected homologs, an unrooted maximum likelihood phylogeny was computed using LG+G4 as model for protein evolution (chosen according to BIC). 100 bootstrap replicates were computed; only bootstrap values ≥50 are shown and bootstrap values of 100 are depicted by a filled dot. Colored font and dots correspond to the support recovered for the higher-order clades labeled on the right of the phylogenies.

The CCR homologs were distributed over several major clades. This included the CCRL/DRL-like sequences described as TETRAKETIDE α-PYRONE REDUCTASE (TKPR) by Grienenberger et al. (2010), which is an important enzyme acting in the production of sporopollenin. We recovered a well-supported clade of TKPR1 homologs (bootstrap support of 84) and fully supported clade of TKPR2 homologs. Both clades of TKPRs contained homologs from across the diversity of land plants, bolstering the idea that TKPR1 and TKPR2 split early during plant evolution (Grienenberger et al., 2010)—before the most recent common ancestor of land plants came about. Our domain structure analyses found that TKPR1 possessed the NAD-dependent epimerase/dehydratase (IPR001509) and NAD(P)-binding domain superfamily (IPR036291) domains, which appear to be present in most sequences included in the phylogeny as well as the Tetraketide alpha-pyrone reductase 1 (IPR033267) domain (Figure S4). In contrast, TKPR2 only encoded the first two domains, which is more similar to what is found in the CCR clade.

The *bona fide* CCRs and CCR-likes were spread out over two clades. These two clades were nested in a weakly-supported monophylum (bootstrap support of 53), which was sub-divided into four medium to fully supported clades. A fully supported clade of CCR-likes (including AT4G30470 and AT2G23910) included sequences from across embryophytes; we coined this monophylum CRL-A. *At*CCRL1 and *At*CCRL2 appear to be co-orthologs to one sequence in the Brassicaceae *Capsella grandiflora* (0380s0077.1.p), thus our data suggest a limited distribution of direct orthologs to CCRL1 and CCRL2. Yet the two sequences together fall into a large clade, here coined CRL-B (bootstrap support 73), that contained sequences from all major lineages of tracheophytes. Another medium-supported clade (bootstrap support 73) included the *bona fide* CCRs, CCR1 and CCR2, of *Arabidopsis.* The duplication that resulted in these two CCRs occurred earliest in the common ancestor of all rosids and latest in the common ancestor of Brassicaceae, yet the CCR clade included homologs from across tracheophytes. Many of these lineages appear to have expanded their own repertoire from one CCR1/2 homolog that was present in the last common ancestor of tracheophytes. Interestingly, a clade of divergent monocot CCRs display several replacements in key amino acids involved in the binding of the substrate’s phenolic ring, in particular a replacement of non-polar aliphatic Ile to aromatic Tyr/Phe (Figure S5), which might reflect a difference in substrate affinity. The catalytic triad SYK (Figure S5; Pan et al. 2014) is required for enzymatic activity and is overall conserved across CCR/DFR-likes and FLDHs. Interestingly, several CRL-As possess non-conservative amino acid replacements from large phenolic (Tyr/Phe) to smaller (His, Leu, Ser, Gly) amino acids, which might suggest divergent substrate affinities for CRL-As. This is consistent with the domain structure of many CRL-A sequences, which often lack the NAD-dependent epimerase/dehydratase (IPR001509) domain, but possess additional domains such as 3-β-hydroxysteroid dehydrogenase/isomerase (IPR002225) or match an additional NAD(P)-binding domain (IPR016040; Figure S4). This pattern is only occasionally occurring in sequences from the CCR or other CRL clades.

Finally, there is a third clade with a bootstrap support of 69 that included tracheophyte sequences (forming a sub-clade with a bootstrap value of 90) and a single *Marchantia polymorpha* homolog; we coined this clade CRL-C. Altogether, this suggests that the LCA of all land plants had two homologs of CCRs/CRLs: one CRL-A and one CRL-B, CRL-C or CCR homolog. In vascular plants, duplications have resulted in sub-clades of the CRL-B/CRL-C/CCR homologs.

Within the larger clade that encases the DFRs, DFRLs, CCRs and CCRLs (“Chloroplastida CCR/DFR-likes”; bootstrap 86), one supported clade of Zygnematophyceae (bootstrap support 77) and one supported clade of chlorophyte and streptophyte algae (bootstrap support 76) exists. This points to a distinct DFR/DFRL/CCR/CCRL clade that arose in the ancestor of Zygnematophyceae, yet its placement within the phylogeny other than it belonging to the larger DFR/DFRL/CCR/CCRL clade is uncertain. The divergent pattern of amino acids, which perform substrate and cofactor binding in land plants, suggest that these algal homologs might vary in the substrate and enzymatic activity compared to plant CCR/DFRs (Figure S5) It is, however, certain that within land plants, a pronounced radiation of CCRs occurred.

### CADs are present across the green lineage

The second reduction step of the activated acyl-group of the phenylpropanoid backbone and one of the last steps in lignin biosynthesis is the production of phenylpropanoid-derived alcohols from the corresponding aldehydes. An example is the conversion of *p*-coumaroyl aldehyde into *p*-coumaryl alcohol (Kim et al., 2004; Pan et al., 2014). The required reduction is catalyzed by CAD (Figure 1), which is the rate determining enzyme by which lignin is produced (Gross et al., 1973; Mansell et al., 1974). In *Arabidopsis thaliana*, there are at least eleven enzymes belonging to the CAD family. Enzymes of the CAD family have been divided into five major groups, of which group IV was described as monocot-specific (Saballos et al., 2009). In our previous studies many of the chlorophyte and streptophyte potential CAD homologs, identified mostly from transcriptomes and few genomes of algae, were described as CAD-like or CAD group II/III-affiliated (de Vries et al. 2017; de Vries et al. 2020). Sequences clustering with those of CAD group II have been characterized as **s**inapyl **a**lcohol **d**ehydrogenase (SAD) or show predicted structural similarity to SAD enzymes (Guo et al. 2010; de Vries et al. 2017). Additionally, some SADs appear involved not in the synthesis of lignin but defense compounds such as lignans (Suzuki and Umezawa, 2007; Guo et al., 2010; Saleem et al., 2010; Barakate et al., 2011), and it may thus be that CAD group II is functionally versatile.

Here, we used the 11 canonical CAD sequences to understand the diversity in CAD homologs across streptophytes. This includes also CAD group II sequences, for which homologs in other species may have other substrate specificities and thus are involved in different steps of the phenylpropanoid pathway (Barakat et al., 2009; Guo et al., 2010). We computed a phylogeny of CAD homologs (Figure 5) detected in phylodiverse Chloroplastida. While the resolution of the backbone is weak, we recovered all five CAD groups defined by Saballos et al. (2009). All the CAD groups were resolved as land plant-specific clades of CAD homologs with robust support. Each clade contained a varying set of major land plant lineages (described below); the clades of putative streptophyte algal CAD homologs contained both fewer proteins and fall in-between the five CAD-groups. Hence, this more phylodiverse dataset tells a more complicated evolutionary history for CAD homologs than the less-diverse data from de Vries et al. (2017).

**Figure 5.**
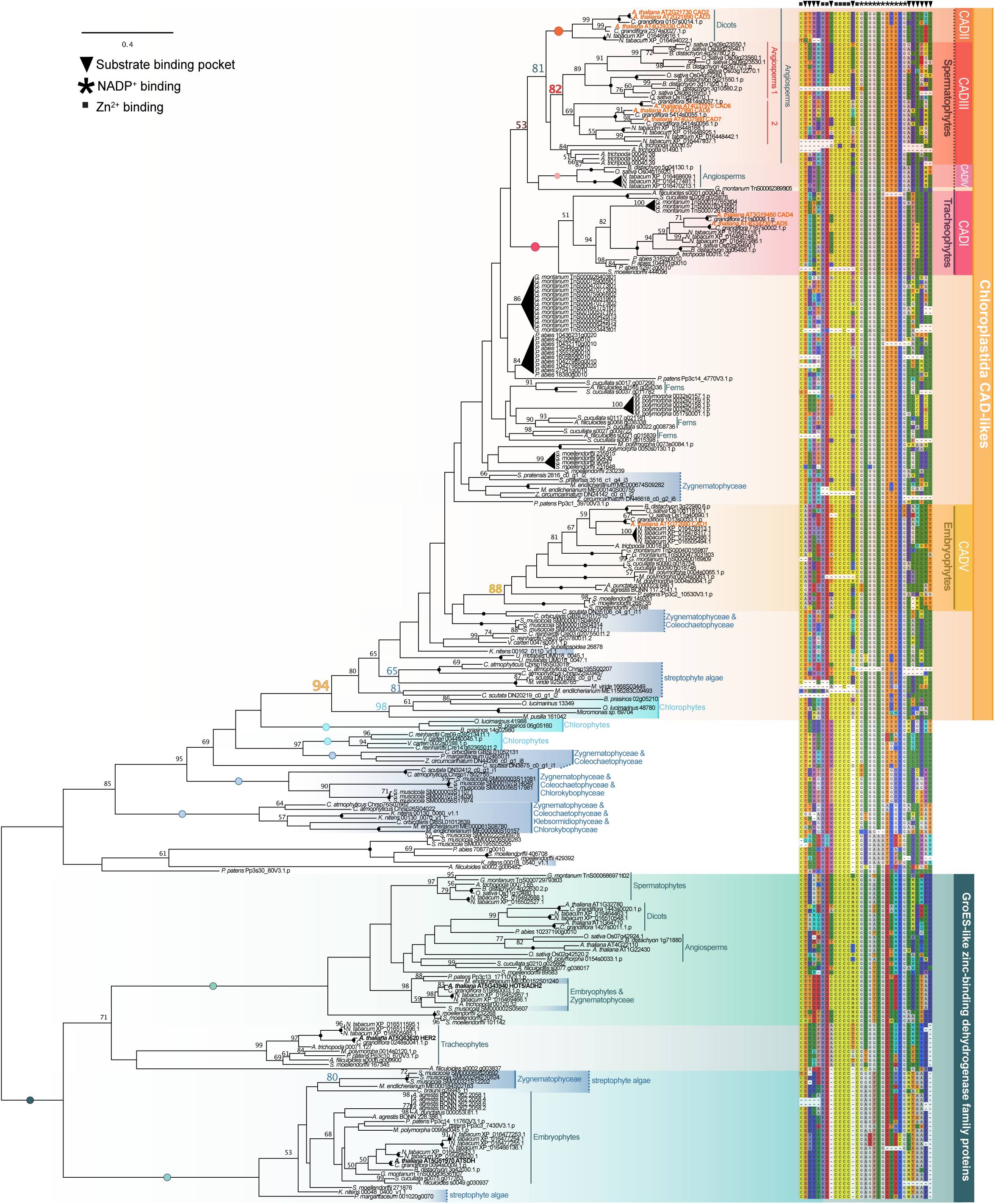
Phylogenetic analysis highlights CAD candidates across Chloroplastida. CAD homologs were sampled from protein data from fifteen land plant, seven streptophyte algal, and five chlorophyte algal genomes as well as sequences found in the transcriptomes of *Spirogyra pratensis* (de Vries et al., 2020), *Coleochaete scutata* and *Zygnema circumcarinatum* (de Vries et al., 2018), and *C. orbicularis* (Ju et al., 2015). From all detected homologs, an unrooted maximum likelihood phylogeny was computed using LG+I+G4 as model for protein evolution (chosen according to BIC). 100 bootstrap replicates were computed; only bootstrap values ≥50 are shown and bootstrap values of 100 are depicted by a filled dot. Colored font and dots correspond to the support recovered for the higher-order clades labeled on the right of the phylogenies. The five groups of CADs were named in accordance with Saballos et al. (2009). Next to the sequence labels residues from the binding pocket, NADP^+^- and Zn^2+^ binding are shown—based on Youn et al. (2016).

Our data suggest that the common ancestor of Zygnematophyceae and land plants may have possessed two CAD-like genes, which was followed by lineage-specific radiations. While it appears—based on the overall topology of the tree—appealing to suggest that one gene gave rise to CAD-group V and the other ancestral gene was the basis of CAD-group I to IV, the low statistical support for the backbone of the phylogeny does not allow to confirm such hypothesis (Figure 5). We can infer that the earliest land plants likely inherited a few (or just one) CAD homolog from their algal progenitors. Most of the radiation of CADs has occurred in plants dwelling on land. Of the canonical CAD group—and based on those part of the topology with good bootstrap support—CAD-group V (containing *At*CAD1) is the only group present in all major land plant lineages (Figure 5). CAD-group I was likely present in the LCA of tracheophytes, as it includes sequences from angiosperms, gymnosperms, ferns, and the lycophyte *Selaginella moellendorffii.* CAD-groups II, III, and IV include only angiosperm sequences—but note that with very weak bootstrap support (53) a sequence from the gymnosperm *Gnetum montanum* associates with the group CADII/III/IV; likely, expansion resulted in the ancestral gene of CAD-group II and III, which diverged into CAD group III genes in angiosperms and after another expansion CAD-group II originated in the LCA of dicots. This is in contrast to de Vries et al. (2017), where the streptophyte algal CAD-like sequences were clustering with CAD group II/III sequences, but resembles the placement of transcriptomic CAD-like sequences from *Spirogyra pratensis* and *Mougotia* sp. in an already more diverse phylogenetic analysis (de Vries et al., 2020). This is a clear case where including a larger diversity of streptophyte sequences to the analysis enables us to better understand the complexity of the evolution of highly radiated gene families. An analysis of the residues salient to CAD function showed a general conservation of residues involved in zinc and NADP^+^ binding across Chloroplastida CAD-like sequences (cf. Youn et al. 2006), whereas the residues in the binding pocket are generally less conserved in these sequences (Figure 5). However, within a canonical CAD-groups or a CAD-like clade we see a general conservation of residues in the binding pocket. The binding pockets of each CAD-like clade appear different than those of the canonical CAD-groups. Domain analyses suggest that some of the CAD-like sequences of streptophyte algae do not encode all of the five domains present in most canonical CAD sequences, yet many of the Zygnematophyceaen CAD-like sequences encode all of these five domains. Similar patterns emerge for other CAD-like sequences (Figure S6).

All CAD groups are shaped by multiple lineage-specific duplications and losses. This hampers the inference of function and substrate specificity of the diverse CAD-like sequences. Additionally, several lineages have originated a variety of CAD homologs that are not yet designated to previous groups. In the absence of functional data, we however will not give them a group designation but rather designate them as lineage-specific CAD-homologs of unknown SAD or CAD function. CAD-like homologs found in streptophyte algae show similar functional residues to other CAD-likes of land plants (not included in any of the five clades of canonical CADs), including at the binding pockets—the pattern of residues is similar to what is observed for other land plant sequences in-between the *bona fide* CAD groups. The *bona fide* CAD groups showed a more homogeneous pattern of functional residues.

Overall, both the topology of the phylogenetic tree and the conservation of key residues point to (i) a deep evolutionary origin of CAD homologs and (ii) independent radiations of CADs—not only in land plants but also in streptophyte algae.

### Massive independent radiations of acyltransferases and scattered candidates in streptophyte algae

A versatile group of enzymes that are important for the processes leading up to the lignins but also compounds with antioxidant and antimicrobial properties are the BAHD acyltransferases (named after the first enzymes characterized for this family BEAT, AHCT, HCBT, and DAT; see also D’Auria, 2006). Most prominent among these are the versatile hydroxycinnamoyl-CoA shikimate/quinate hydroxycinnamoyltransferases (HCT) (Eudes et al., 2016). Recently, Kriegshauser et al. (2020) reported on the functional conservation of the HCT homologs found in bryophytes with those of seed plants.

We computed a phylogeny including phylodiverse acyltransferases (Figure S7). Overall, the topology of the tree corroborates the findings of Kriegshauser et al. (2020) that a clear clade of HCT proteins first emerged in land plants—streptophyte algal sequences were few, divergent from HCT, and scattered over the tree without clear affinity to characterized acyltransferases. Without functional analyses, there is no solid foundation for predicting their function—making them exciting candidates for future studies. What our data, however, clearly reveal is that the last common ancestor of land plants likely had an expanded repertoire of acyltransferases that further diversified during the radiation of plants on land.

### A clear clade of C3Hs is limited to land plants

At several steps of the phenylpropanoid pathway, enzymes belonging to the cytochrome P450 family CYP98A, within the large CYP71 clan (Nelson and Werck-Reichhart, 2011), carry out hydroxylations of *p*-coumarate-derived compounds (such as *p*-coumaroyl esters; for function of C3H see Figure 1). In land plants, these hydroxylations are important for the production of lignins, lignans, volatile phenylpropanoids, coumarins, and many more phenylpropanoid-derived compounds. Previously, de Vries et al. (2017) described the detection of C3H in all land plants and one putative C3H ortholog in *Klebsormidium nitens*. Now, with genomic gaps in the streptophyte tree of life filled, we revisited the distribution of C3H.

The number C3H homologs detected in genomes of 15 land plants, 7 streptophyte algae, and five chlorophytes and in the transcriptomes of *Spirogyra pratensis* (de Vries et al., 2020), *Zygnema circumcarinatum* (de Vries et al., 2018), and *Coleochaete orbicularis* (Ju et al., 2015) varied strongly between lineages. When sampling the sequences via BLAST (with AT2G40890 as query sequence), we thus included either (a) all sequences that had a bit score of at least 100, or (b) the top five hits. We aligned all sequences, cropped them to the alignable region and computed a maximum likelihood phylogeny (Figure 6).

**Figure 6.**
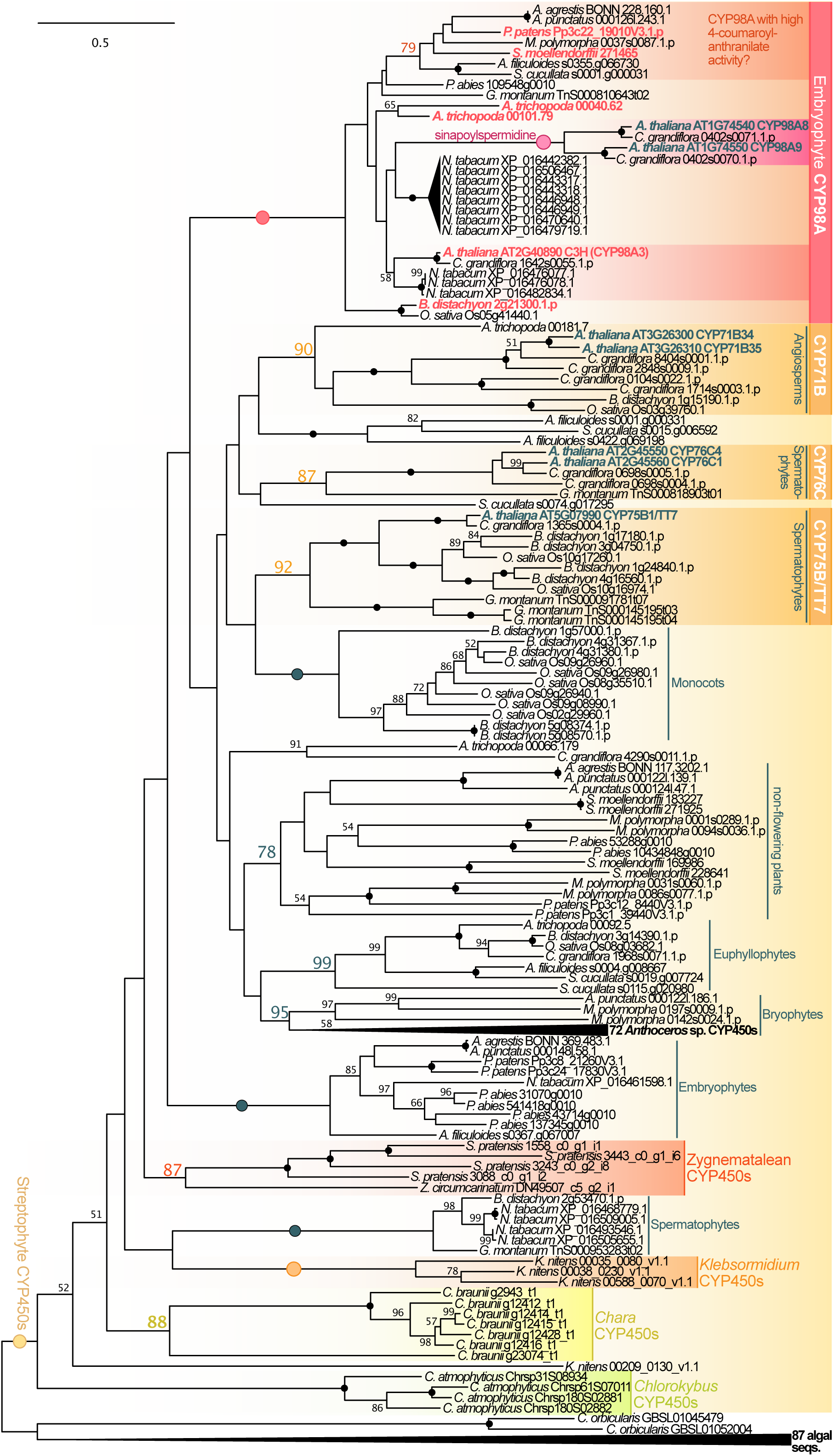
A clade of C3H orthologs originated at the base of land plants. C3H homologs were sampled from protein data of genomes of fifteen land plants, seven streptophyte algae, and five chlorophytes; additionally, sequences found in the transcriptomes of *Spirogyra pratensis* (de Vries et al., 2020), *Zygnema circumcarinatum* (de Vries et al., 2018), and *Coleochaete orbicularis* (Ju et al., 2015) were included. For downstream analyses, we used either (a) all sequences that had a bit score of at least 100 or (b) the top five hits. We aligned all sequences, cropped them to the alignable region and computed an unrooted maximum likelihood phylogeny was computed using LG+F+I+G4 as model for protein evolution (chosen according to BIC). 100 bootstrap replicates were computed; only bootstrap values ≥50 are shown and bootstrap values of 100 are depicted by a filled dot. Colored font and dots correspond to the support recovered for the higher-order clades labeled on the right of the phylogenies. Two large clades that contained only (a) *Anthoceros* and (b) chlorophyte and streptophyte algal sequences were collapsed; the full tree is shown in Figure S12.

A clade of CYP98A included sequences from all major lineages of land plants. This suggests that at least one CYP98A sequence was present in the LCA of all land plants. Based on the lineages included here, it appears that from this single copy gene, radiations occurred in the dicot lineages and *Amborella trichocarpa*. A single copy remained in the bryophytes, lycophytes, ferns, gymnosperms, and monocots. The clade of CYP98A8 and CYP98A9 was in our dataset limited to the Brassicaceae *Arabidopsis thaliana* and *Capsella grandiflora* (bootstrap value 100). These two enzymes function in a route derived from the phenylpropanoid pathway and are involved in the formation of *N*^1^,*N*^5^-di(hydroxyferuloyl)-*N*^10^-sinapoylspermidine (Matsuno et al., 2009). The CYP98A8/9 clade falls into the larger CYP98A clade (bootstrap support 100) together with the C3H sequences, suggesting that they are the closest paralogs of C3H in *Arabidopsis thaliana* and *Capsella grandiflora.* Analyses of their substrate recognition sites (SRS; cf. Rupasinghe et al. 2003) support the divergent functional roles between CYP98A8/9 and canonical C3H. In particular, the first two SRS (SRS1 and SRS2) show various distinct amino acid differences between the CYP98A8/9 clade and the C3H clade (Figure S8). SRS1 and 2 are predicted to be involved in binding of the substrate tails, which exhibit strong variation in size (Rupasinghe et al. 2003), and thus may be critical for the substrate specificity of these paralogs. Indeed, SRS1 and 2 are the two SRSs showing the strongest variation across the entire phylogeny, including also other CYP450 subfamilies (Figure 6), corroborating this hypothesis.

We recovered additional land plant-specific clades of CYP450 enzymes (Figure 6) such as one containing TRANPARENT TESTA7 (TT7)-like sequences (bootstrap support 92); TT7 is a cytochrome P450 75B enzyme that is involved in flavonoid biosynthesis (Tanaka et al., 1997; Schoenbohm et al., 2000). The BLAST search used for sampling C3H homologs further recovered *At*CYP71B and *At*CYP76C members. In our phylogenetic analysis, we inferred that *At*CYP71B34 and *At*CYP71B35 were likely born out of an *Arabidopsis*-specific duplication, while CYP71B enzymes in general are present across angiosperms. In contrast, *At*CYP76C1 and *At*CYP76C4 appear to have originated prior to the split of *Arabidopsis thaliana* and *Capsella grandiflora*; a CYP76C4 ortholog appears to have been lost in the latter plant species. The CYP76C clade may also be represented in other species outside of angiosperms, because we find a sequence from *Gnetum montanum* clustering with these sequences with a bootstrap support of 87. The rather long branch warrants attention and would require a more CYP76C focused phylogenetic analysis, which is not the point of this paper.

The algal sequences showed strong divergence to the C3Hs, forming only a larger (fully supported) streptophyte-specific clade with all the recovered and functionally diverse CYP450 enzymes (Figure 6). Domain structures of these streptophyte algal sequences is the same to what is observed for the C3H homologs, but is in general conserved across the phylogeny independent of the CYP450 subfamily assignment (Figure S9). Only few scattered exceptions occur. Given the low support of most of the tree backbone, the role(s) of the streptophyte algal homologs detected here remains elusive. Most of the streptophyte algae show independent radiations of their CYP450 enzymes complicating functional predictions even further. Thus, while there are interesting CYP450 candidates in streptophyte algae, a clear C3H clade likely first arose early during the evolution of embryophytes.

### Monoacylglycerol lipases: multiple early radiations, independent subfunctionalization, and the origin of CSE

The conversion of caffeoyl-5-*O*-shikimate to caffeic acid may be a step along the biosynthetic routes that lead to the production of G- and S-lignins in certain vascular plants (Figure 1). The enzyme responsible for this step is caffeoyl-5-*O*-shikimate esterase (CSE). CSE converts caffeoyl-5-*O*-shikimate to caffeic acid and was hypothesized to act together with 4CL/ACOS5 to circumvent the catalysis of caffeoyl-5-*O*-shikimate to caffeoyl-CoA via HCT (Vanholme et al. 2013). The latter pathway was proposed for tobacco by Hoffmann et al. (2003) and confirmed for *Arabidopsis thaliana in vitro* by Vanholme and colleagues (2013). Yet, based on *cse* mutants in *Arabidopsis thaliana*, they suggested that synthesis of caffeoyl-CoA is more likely to occur via CSE and 4CL/ACOS5 than directly from caffeoyl shikimate by HCT *in planta.* That said, in the model grasses *Brachypodium distachyon* and *Zea mays* no CSE orthologs are present and crude extracts from these species show little signs for the characteristic esterase activity (Ha et al., 2016). On the other hand, non-vascular plants such as the model system *Physcomitrium patens* possess homologs of these enzymes (Renault et al., 2017a), which suggests a secondary loss of CSE in the respective monocots. CSE belongs to the family of putative monoacylglycerol lipases (MAGL). MAGLs are found across eukaryotes and functional analyses in human, yeast and *Arabidopsis* have shown that they possess monoacylglycerol lipase activity (Labar et al., 2010; Aschauer et al., 2016; Kim et al., 2016). In contrast to other MAGLs of *Arabidopsis thaliana, At*CSE (MAGL3) was found to exhibit no hydrolytic activity on monoacylglycerols (MAGs) as a substrate (Kim et al., 2016)—which applies to other enzymes of *Arabidopsis thaliana* that belong to the family of MAGLs, too. Indeed, out of the 16 MAGLs that Kim and colleagues (2016) tested, only MAGL6 and 8 showed high activity on MAG as substrate. Given the functional diversity in MAGLs (Kim et al. 2016) and the unequal distribution of caffeoyl-5-*O*-shikimate across embryophytes, functional analyses are required to fully understand how easily MAGLs can lose or gain their MAGL activity. Yet, phylogenetic analyses can pinpoint the diversity of the family across the green lineage.

Here we use phylogenetic analysis to pinpoint the distributions of the diverse MAGL families, including CSE across the green lineage. In total, we recovered all 16 MAGL sequences of *Arabidopsis thaliana* in the similarity search; using maximum likelihood phylogenetics, we recovered clades for all the 16 MAGLs; some MAGL clades are widely distributed throughout streptophytes, while others appear to have originated in embryophytes, where they have again undergone lineage-specific expansions. Essentially, we recovered two large clades: one restricted to streptophytes, containing homologs of MAGL2, 4 and 13; the other has representation in chlorophytes as well and includes homologs of MAGL1, 3, 5, 6, 7, 8, 9, 10, 11, 12, 14, 15 and 16 (Figure 7).

**Figure 7.**
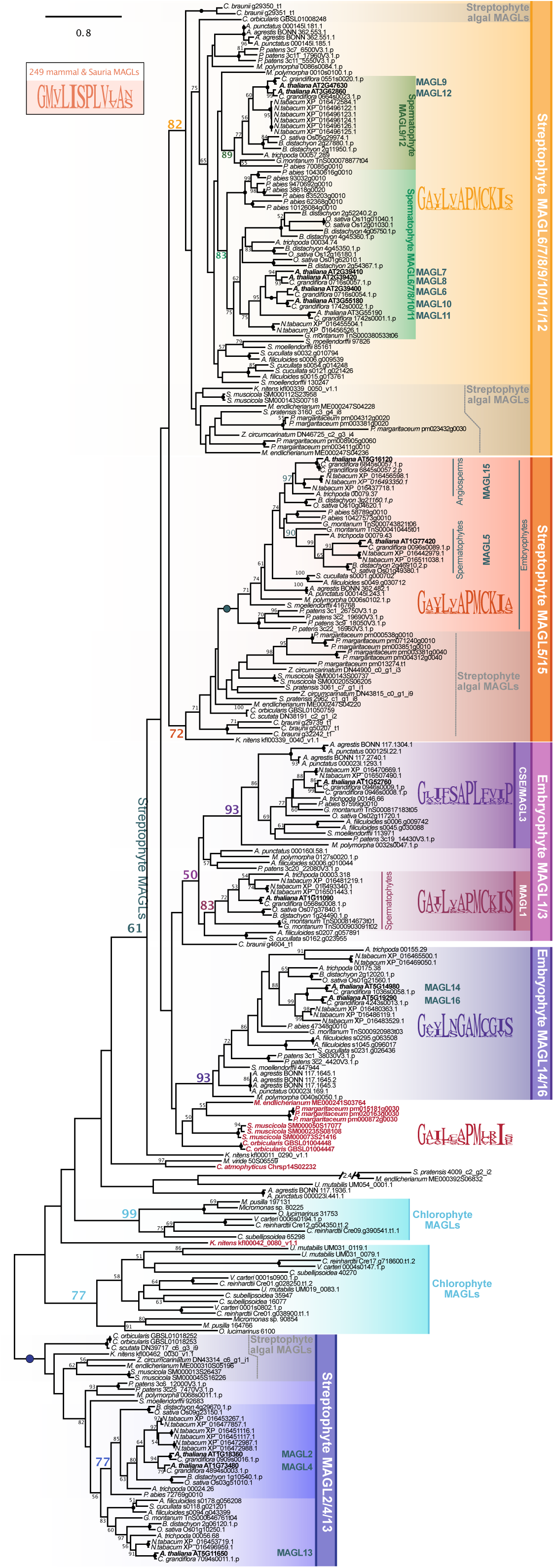
The occurrence of MAGLs across diverse Streptophyta and a phylogenetic framework for the deep evolutionary roots of CSE. MAGL/CSE homologs were sampled from protein data from 15 land plant, seven streptophyte algal, and five chlorophyte algal genomes as well as sequences found in the transcriptomes of *Spirogyra pratensis* (de Vries et al., 2020), *Coleochaete scutata* and *Zygnema circumcarinatum* (de Vries et al., 2018), and *Coleochaete orbicularis* (Ju et al., 2015). From all detected homologs, an unrooted maximum likelihood phylogeny was computed using LG+I+G4 as model for protein evolution (chosen according to BIC). 100 bootstrap replicates were computed; only bootstrap values ≥50 are shown and bootstrap values of 100 are depicted by a filled dot. Colored font and dots correspond to the support recovered for the higher-order clades labeled on the right of the phylogenies. Purple font highlights those streptophyte algal sequences that share the conserved alpha helix cap domain with CSE. Logos are based on a motif (amino acids 132-141 in MAGL6 and 167-176 in CSE/MAGL3) that is situated in a region likely involved in substrate binding based on the crystal structure of human MAGL.

Focusing on the MAGL2/4/13 clade first, we observe that MAGL13 has representation in angiosperms, gymnosperms and ferns, suggesting its origin to be in the LCA of tracheophytes, while MAGL2 and 4 came from a duplication event before the split between *Arabidopsis* and *Capsella.* However, MAGL2/4 orthologs are present in other species including *Picea abies,* suggesting that the common ancestor of seed plants possessed a *MAGL2/4-like* and a *MAGL13* gene. Forming a clade with MAGL2/4/13 are lycophyte, bryophyte and streptophyte algal sequences, which in general branch in an order expected based on their species phylogeny (although within species duplication events have occurred). This suggests that already at the base of streptophytes a *MAGL2/4/13-like* gene was present.

In the second large clade that includes also chlorophyte sequences, we find the clade containing the CSE/MAGL3 orthologs. This clade includes sequences from both vascular and non-vascular plants, pointing to an origin of CSE in the last common ancestor of land plants. This adds support for a secondary loss in those monocots without a CSE ortholog. Despite the origin of CSE in the common ancestor of land plants and a clear CSE ortholog in *Physcomitrium patens* (3c19_14430V3.1.p), the substrate of CSE, caffeoyl-5-*O*-shikimate, was not detected in crude extracts of the moss (Renault et al., 2017a). The HCT-based reaction leading to caffeoyl-CoA has been confirmed *in vitro* using moss HCT (Kriegshauser et al. 2021). Hence, the CSE homologs of *Physcomitrium patens* may have another function. Indeed, the atypical function of *At*CSE, together with the lack of other MAGL family members, to act on MAGs (Kim et al. 2016), suggests that the functional spectrum of the MAGL family is not very limited in land plants.

Members of the MAGL family share several conserved motifs across diverse eukaryotes. One such motif (amino acid positions 132-141 in MAGL6 and 167-176 in CSE/MAGL3) is situated in a region likely involved in substrate binding based on the crystal structure of human MAGL (Labar et al., 2010). Within this motif a leucine in position number four is found in a diverse set of 249 mammal and Sauria MAGLs investigated here (Figure 7, inset) and most plant MAGLS including MAGL6 and 8. This is followed by another hydrophobic amino acid (I in mammals/Sauria and V or L in most plant MAGLs). It is striking that exactly these highly conserved amino acids are changed to a phenylalanine and a serine in the Arabidopsis CSE and some homologs from other species. These changes from two very hydrophobic amino acids to an aromatic and a hydrophilic one could be one of the reasons for a change in substrate specificity from a substrate with a hydrophobic acyl chain to a more hydrophilic substrate with aromatic properties. Based on this hypothesis, CSEs would be restricted to some of the members of this clade that, however, stem from across the diversity of land plants.

MAGL1 appears to have originated prior to the split of angio- and gymnosperms, while specific MAGL14 and 16 orthologs likely arose after the split of asterids and rosids, but a MAGL14/16 ortholog was likely present in the LCA of land plants. MAGL15, like MAGL1, originated prior to the split of gymno- and angiosperms, and MAGL5 appeared to come from a duplication of MAGL15 later on possibly in the ancestors of dicots. Interestingly, a *MAGL5/15-like* sequence was already encoded in the genome of the ancestor of streptophytes. The same is true for a *MAGL6/7/8/9/10/11/12-like* gene, which similar to the *MAGL5/15-like* genes shows independent paths of radiation in streptophyte algae and land plants. MAGL6,7,8,10 and 11 are only present in the here included Brassicaceae, while a *MAGL6/7/8/10/11-like* gene was already present in the common ancestor of spermatophytes. The same evolutionary history describes the scenario under which MAGL9 and 12 originated.

Finding the MAGL6/7/8/10/11 subclade specifically expanded in seed plants is noteworthy. At least *At*MAGL8 localizes to lipid droplets (Kim et al., 2016), which are structures found in various photosynthetic eukaryotes but are well-known from seeds. Thus, expansion of this clade might be a read-out of spermatophyte-specific additions to the ancient set of proteins relevant to LD formation and function (see de Vries and Ischebeck, 2020).

All in all, the MAGLs have experienced an early radiation in streptophytes. Given that even the Arabidopsis MAGLs without detectable activity on MAG (see Kim et al., 2016) do not form one single monophylum, it is conceivable that subfunctionalization of members of the MAGL family occurred multiple times independently. This may likewise be true for all independent expansions of MAGL-encoding genes observed in any other species included here. The versatility in functional evolution of MAGLs makes it difficult to make robust predictions of putative MAGL functions.

### COMT: convergence and complexity

In angiosperms, ferulate 5-hydroxylase (F5H) and caffeate *O*-methyltransferase (COMT) carry out important catalytic steps along the route from *p*-coumaroyl-CoA to S-lignin. COMT catalyzes the methylation of caffeic acid or 5-hydroxyferulic acid, the product formed by F5H. Like C3H and C4H, F5H belongs to the large CYP450 clan 71 (Nelson and Werck-Reichhart, 2011). The function of F5H evolved at least twice in P450 enzymes, once in the ancestor of angiosperms and once in the ancestor of lycophytes (Weng et al., 2008; Weng and Chapple, 2010).

For angiosperm F5H, no clear putative orthologs were found outside of flowering plants and likewise no clear orthologs were found for the lycophyte F5H (i.e. “SmF5H”), which forms a separate clade from the angiosperm F5H sequences (Figure S10). This corroborates previous results (de Vries et al., 2017) and is in agreement with the hypothesis that F5H function evolved at least twice in the evolution of land plants (Weng et al., 2008). Additionally, the average pairwise identity of the F5H homologs was quite low (18.5%)— hampering robust phylogenetic analyses. We thus did not further delve into the evolution of F5H. COMT however caught our attention.

The lycophyte *Selaginella moellendorffii* not only uses a genetically distant F5H enzyme; the same appears to be true for COMT (Weng et al., 2011). This highlights a promiscuity for substrate specificity and activity in P450 enzymes that is yet to be discovered and mere orthology analyses can only go so far as to discover putative candidates. Using phylogenetics, we explored the diversity of methyltransferases by screening for sequences homologous to COMT/OMT1 of *Arabidopsis thaliana* across our phylodiverse dataset. This approach identified not only clear orthologs but can also serve as a backbone to map relevant mutations facilitating in functionally convergence in this group of enzymes and by that may highlight possible candidates for *in vivo* and *in vitro* studies.

For most clades of land plant methyltransferases, based on the here recovered topology, predicting a putative function was not straightforward. This applied even more so to the homologs of COMT/OMT found in chlorophyte and streptophyte algae. We recovered a clade of methyltransferases that included chlorophyte and streptophyte algae as well as diverse land plant sequences (coined ‘Chloroplastida OMT’ in Figure 8); among these clustered *Arabidopsis thaliana* proteins such as COMT, indole glucosinolate methyltransferases (IGMT), and nicotinate *N*-methyltransferase (NANMT; see Li et al., 2017)—hence different methyltransferases that act on a range of aromatic compounds. What this means for the presence of a putative COMT in algae is obscure. However, it corroborates the previously observed patchy detection of OMT1 across the green lineage based on a reciprocal BLASTp searches (de Vries et al., 2017). Clear orthologs of *At*COMT were only detected for a few angiosperms, notably not including any monocot sequence that we included in our dataset. Of all methyltransferases in our dataset, only NANMT formed a clade of clear orthologs that included more than one major lineage of land plants by encasing sequences from angiosperms and *Picea abies* (bootstrap support 87). All other orthogroups appear, like COMT, to be restricted to only a few of the included angiosperm lineages.

**Figure 8.**
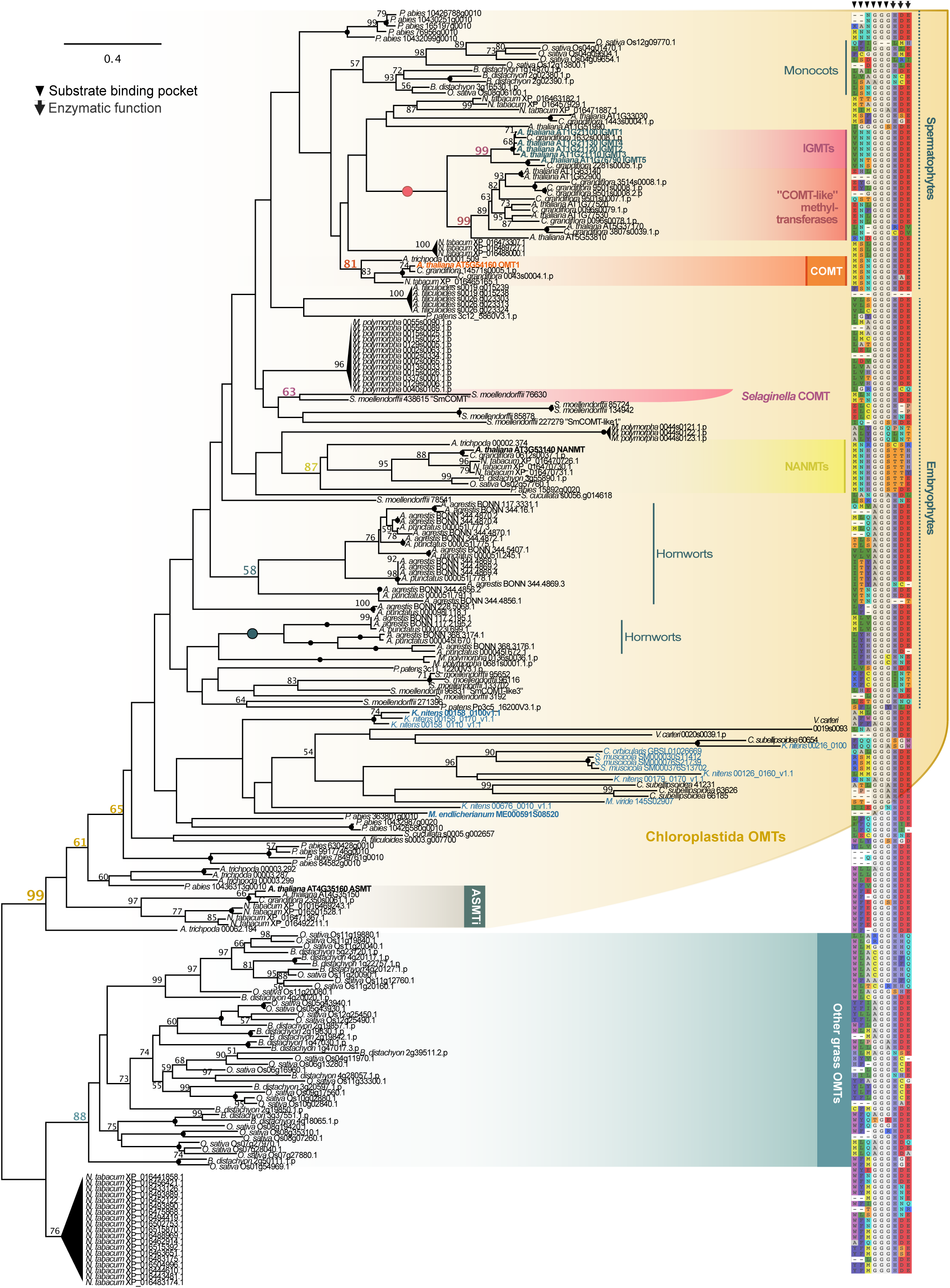
Low resolution on the complex evolutionary history of COMT. We explored the diversity of methyltransferases by screening for sequences homologous to Arabidopsis COMT/OMT1 across genome data from fifteen land plant, seven streptophyte algae, and five chlorophytes; additionally, we included sequences found in the transcriptomes of *Spirogyra pratensis* (de Vries et al., 2020), *Zygnema circumcarinatum* (de Vries et al., 2018), and *Coleochaete orbicularis* (Ju et al., 2015). From all detected homologs, an unrooted maximum likelihood phylogeny of 226 sequences was computed using LG+I+G4 as model for protein evolution (chosen according to BIC). 100 bootstrap replicates were computed; only bootstrap values ≥50 are shown and bootstrap values of 100 are depicted by a filled dot. Colored font and dots correspond to the support recovered for the higher-order clades labeled on the right of the phylogenies. Blue font highlights streptophyte algal sequences; bold font pinpoints those, that recovered land plant COMT as closest structural analogs in I-TASSER-based modeling. On the right we show residues important for substrate binding and function of canonical COMT as reported by Louie et al. (2010).

The lycophyte *Selaginella moellendorffii* has a COMT that is distantly related to OMT of angiosperms. It appears to have acquired its OMT activity through convergent evolution and was coined *Sm*COMT (Weng et al., 2011). In agreement with this, *Sm*COMT did not cluster with the *At*OMT1 sequence in our analyses. Instead, it forms its own (weak) clade with only one other sequence from *Selaginella moellendorffii* (bootstrap support 63). The other *Sm*COMT-like sequences (described in Weng et al., 2011) were distributed over the phylogeny and appear to be specific to *Selaginella moellendorffii.* Nonetheless, this pattern highlights a certain versatility in the evolutionary history of substrate specificity of *O*-methyltransferases in land plants. This appears to be only logical, noting the large lineage-specific expansions in the larger clade of *O*-methyltransferases that encompasses all *O-*methyltransferases from *Arabidopsis thaliana*—that is COMT, IGMTs, NANMT and *N*-acetylserotonin *O*-methyltransferase (ASMT; for more on this enzyme see, e.g., Tan et al., 2012; Byeon et al., 2016) (bootstrap-support 99). Within this clade fall also algal sequences from chlorophytes and streptophyte algae. Their position within the clade is undetermined due to low bootstrap support. These sequences appear highly divergent, many of them cluster with rather long branches. Yet, some of the sequences from our previous analysis found a reciprocal BLASTp hit to *At*COMT, including *Klebesormidium nitens* 00158_0100v1.1 that clusters in a fully supported clade of *Klebesormidium* paralogs; these sequences are promising candidates to explore caffeic acid *O-*methyltransferase activity. Indeed, when we modelled the tertiary structure of *Klebesormidium nitens* 00158_0100v1.1 and *Mesotaenium endlicherianum* ME000591S08520 using I-TASSER (Zhang, 2008), we recovered *Medicago sativa* and *Lolium perenne* COMT as its closest structural analogs (1KYZ; Zubieta et al., 2002; 3P9C; Louie et al., 2010; TM-scores 0.865 and 0.956, respectively).

Like the land plant COMTs, also the algal COMT-likes appear to have undergone independent radiations in this large gene family. Given the observed convergent evolution of COMT activity in *Selaginella moellendorffii*, the question of whether there is COMT activity across Streptophyta remains wide open.

To gain a first insight into whether COMT activity can be expected from other streptophyte lineages, we investigated the conservation of residues relevant for the function of COMT, including those that form the substrate binding pocket (Figure 8). The functional residues were identified from COMT of *Lolium perenne* (Louie et al., 2010). Across our phylogeny, these residues differ between the clades of canonical ASMT, NANMT, COMT and IGMT, while they are conserved within them (Figure 8). The binding pocket of *At*OMT1 and its orthologs consist of the amino acid pattern MSNGGG, whereas the pattern for the residues important for the function of the enzyme is HDE. While HDE appears conserved across the majority of sequences analyzed here, independent of the specific function of the enzyme (e.g. ASMT, IGMTs and COMT all have the pattern HDE), the binding pocket is highly variable among the functionally characterized enzymes. This suggests that the reaction-determining residues are those that form the binding pocket and not those that are catalytically important. This seems logical given that all these enzymes catalyze similar types of reactions. The triple G in the binding pocket is also far more conserved across the entire phylogeny, with only few exceptions occurring, while the first three residues are highly variable. Indeed, the COMT-specific MSN motif is not present in the functionally characterized COMT from *Selaginella moellendorfii,* rather it is MTN, which however is a change between similar amino acids (Ser to Thr). Apart from the COMT orthologs and *Sm*COMT sequence no other sequences from any other lineage encode the binding pocket pattern M(S/T)NGGG, suggesting that none has a canonical preference for binding 5-hydroxyconiferalaldehyde. Yet, several homologs—including those of streptophyte algae— would have the ability to catalyze the reaction based on the conservation of the residue pattern HDE. What is more, all the sequences that could not be properly identified as orthologs to ASMT, NANMT, COMT and IGMT (with the exception of *Sm*COMT) show no similarity in their first three residues of the binding pocket to either of these enzyme families. This would suggest that the enzymes from most streptophyte lineages included in this analysis use different substrates than those functionally characterized in *Arabidopsis thaliana*.

### CCoAOMT-like sequences emerged in Phragmoplastophyta

Within the phenylpropanoid pathway, caffeoyl-CoA *O*-methyltransferase (CCoAOMT) and most of its homologs are the enzymes that catalyze the first committed step to many of at least two types of lignin (S- and G-lignin). These enzymes methylate caffeoyl-CoA and thus give rise to feruloyl-CoA (Ye et al., 1994; Ye and Varner, 1995; Martz et al., 1998; Do et al., 2007; Vanholme et al., 2012); in the past, it was also proposed that after the conversion to 5’-hydroxy-feruloyl-CoA, CCoAOMT can methylate this compound to produce Sinapoyl-CoA (Maury et al., 1999; Ferrer et al., 2005). One of the paralogs that exist in *Arabidopsis thaliana* (tapetum-specific *O*-methyltransferase [TSM1], AT1G67990) however, shows activity towards a coniferyl derivative that is formed at *N*^10^ by F5H starting from *N*^1^,*N*^5^,*N*^10^-tris-(hydroxyferuloyl) spermidine (Fellenberg et al., 2009); TSM1 catalyzes the production of *N*^1^,*N*^5^-bis-(hydroyferuloyl)-*N*^10^-synapoylspermidine (Fellenberg et al., 2009). Thus, CCoAOMTs appear to be versatile in their substrate specificity and can act on different steps in the phenylpropanoid pathway.

Here we used phylogenetics to disentangle the distribution of these enzymes across the green lineage. A duplication gave rise to the genes encoding the functionally divergent enzymes *At*CCoAMT and *At*TSM1 (bootstrap support 77; Figure 9). These two methyltransferases are embedded in a larger clade containing the other CCoAOMT enzymes *At*CCoAOMT1, *At*CCoAOMT7 and *At*CCoAOMT-like (AT1G24735). The latter appears to be specific to the Brassicaceae included in this dataset, while homologs of *At*CCoAOMT7 occur across dicots and were detected in *Amborella trichopoda,* but were absent from the included monocots. Only *At*CCoAOMT1 had a wider distribution. Its cluster (bootstrap support 79) contains angiosperms, gymnosperms, lycophytes, ferns and bryophytes, excluding the sequenced hornworts from the genus *Anthoceros* (Figure 9). Assuming a monophyly of Bryophyta (Puttick et al., 2018), this suggests a loss of CCoAOMT1 in at least the sequenced *Anthoceros* species, and that CCoAOMT1 was present in the common ancestor of land plants. Lineage-specific duplications of CCoAOMT1 appear to have happened, indicated by the expansions seen in tobacco, spruce, the lycophyte *Selaginella moellendorffii* and the water fern *Azolla filiculoides.* The expanded repertoire of sequences in monocots and *Gnetum* indicate additional lineage-specific duplications outside of the CCoAOMT clade. The case of TSM1 suggests that neo-functionalization can easily occur within this type of methyltransferases. We noted that the residues involved in substrate binding (Ferrer et al. 2005) are identical in *At*CCoAMT and *At*TSM1 (Figure 9). A possible explanation might be that the make-up of binding pocket allows for a certain versatility in substrates. Given these observations, the paralogs within this and the other CCoAOMT clades cannot be assumed to hold the function of CCoAOMTs. Likewise, it cannot be ruled out that their LCA may have had this function. As sister to the methyltransferase clade, including the CCoAOMT homologs lies a cluster of genes encoding putative candidates for streptophyte algal CCoAOMTs. These were limited to representatives of the two streptophyte algal lineages closest to land plants: the Coleochateophyceae *Coleochaete scutata* and *Coleochaete orbicularis* as well as the Zygnematophyceae *Spirogloea muscicola*. These algal sequences have the same domain structure as the majority of all *S*-adenosyl-*L*-methionine (SAM)-dependent methyltransferases included in the phylogeny (Figure S11). Only single sequences, scattered across the phylogeny and diversity of species included here, vary in their domain structure showing a loss of a domain loss, a gain of an additional domain, or both. The analyses of the specific functional residues gave more insights into the streptophyte algal sequences within the clade of SAM-dependent methyltransferases in the peripheral routes of the phenylpropanoid pathway. These algal sequences maintain the residues involved in ion and cofactor binding, but differ strongly in the substrate binding site (cf. Ferrer et al. 2005; Figure 9).

**Figure 9.**
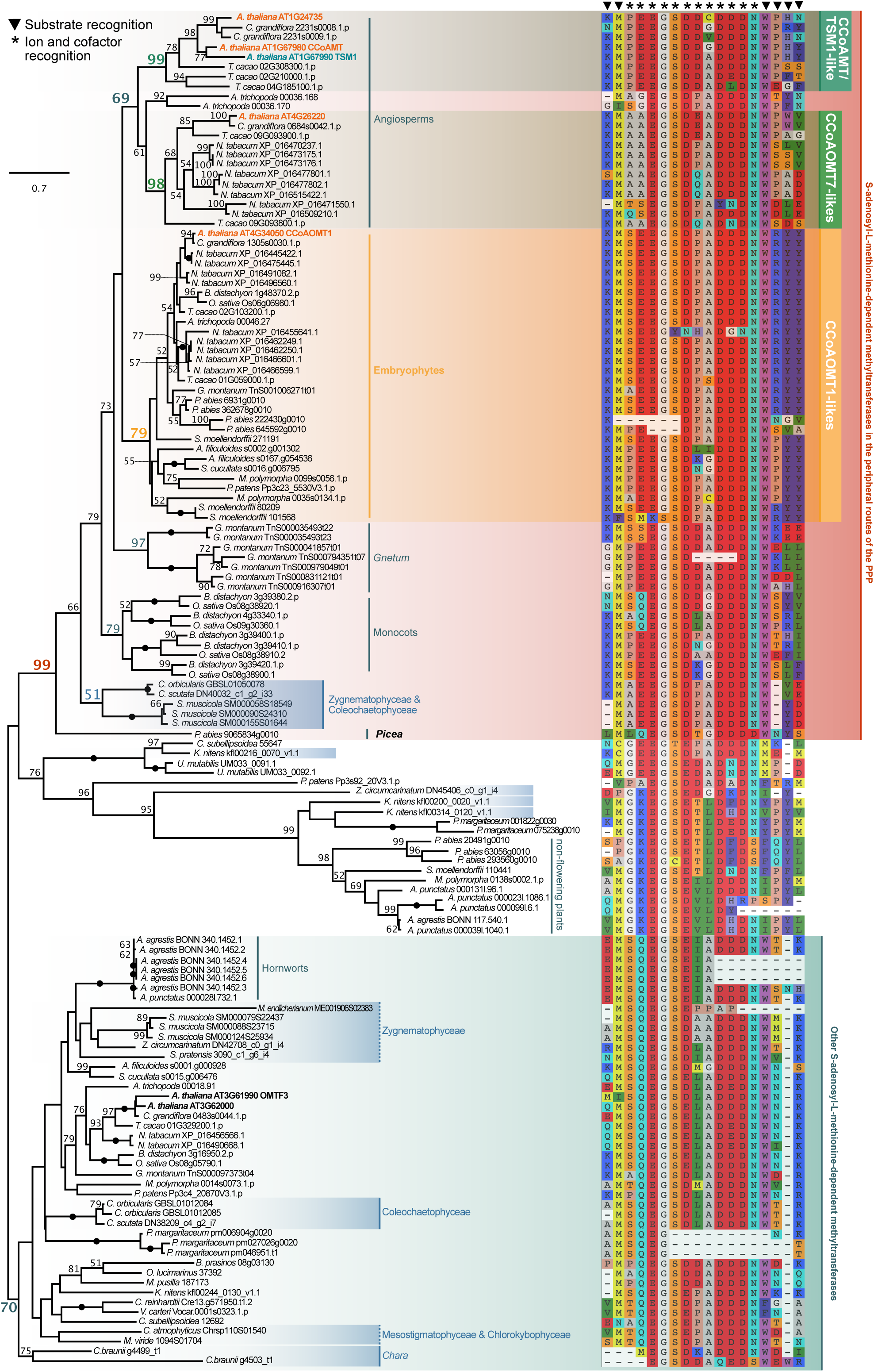
A phylogenetic framework for the evolutionary origin of CCoAOMTs in Phragmoplastophyta. CCoAOMT homologs were sampled from protein data from sixteen land plant, seven streptophyte algal, and five chlorophyte algal genomes as well as sequences found in the transcriptomes of *Spirogyra pratensis* (de Vries et al., 2020), *Coleochaete scutata* and *Zygnema circumcarinatum* (de Vries et al., 2018), and *Coleochaete orbicularis* (Ju et al., 2015). From all detected homologs, an unrooted maximum likelihood phylogeny of 138 sequences was computed using LG+G4 as model for protein evolution (chosen according to BIC). 100 bootstrap replicates were computed; only bootstrap values ≥50 are shown and bootstrap values of 100 are depicted by a filled dot. Colored font and dots correspond to the support recovered for the higher-order clades labeled on the right of the phylogenies. Purple font highlights those streptophyte algal sequences that share the conserved alpha helix cap domain with CSE. The alignment on the right shows functionally characterized sites involved in substrate, ion and co-factor recognition of CCoAOMT (Ferrer et al., 2005).

Altogether, it appears that the family of CCoAOMT-like proteins has its origin in Phragmoplastophyta. Clarifying the function of the putative CCoAOMT-like enzymes in Coleochateophyceae and Zygnematophyceae has the potential to shed light on a synapomorphy with physiological relevance.

## Conclusion

All genes for enzymes that act in early steps in the chassis of the phenylpropanoid pathway investigated here (Figure 1) can be traced back to the LCA of land plants with the exception of COMT (Figure 10); most of these can even be traced back to some ancestor that land plants shared with streptophyte algae. While most of our knowledge on how these genes work comes from angiosperms, this does not capture the sequence diversity in enzymes—and it underpins the versatility in producing specialized metabolites.

**Figure 10.**
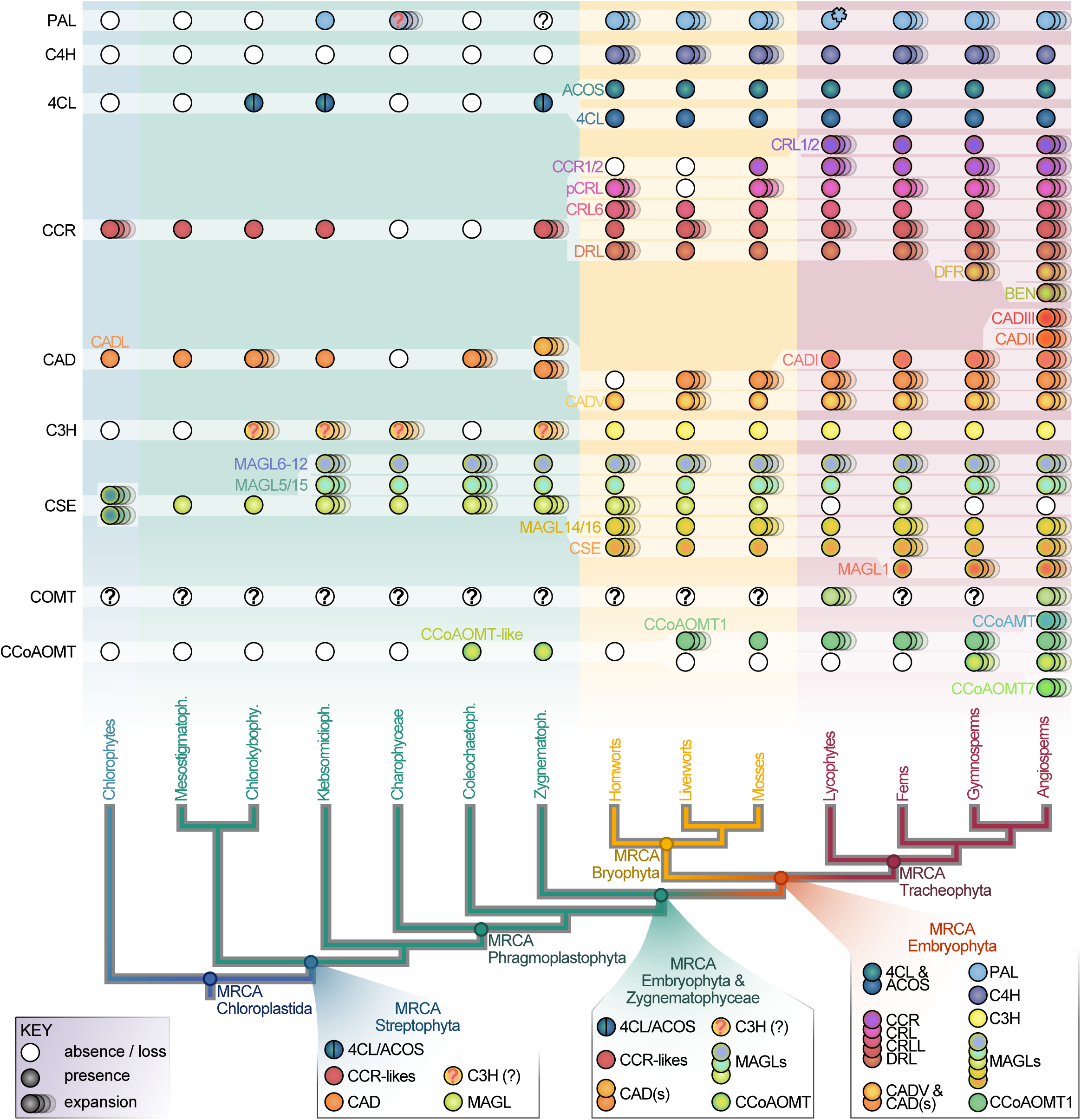
A summary of the proposed evolutionary trajectory of key enzymes in the phenylpropanoid pathway across the green lineage. At the bottom is a cladogram of the green lineage. The most recent common ancestors (MRCA) of Chloroplastida, Streptophyta, Phragmoplastophyta, Bryophyta, Embryophyta and Tracheophyta are indicated at their respective nodes. On top the cladogram is the proposed evolutionary trajectory of the enzyme families PAL, C4H, 4CL, CCR, CAD, C3H, CSE/MAGL (CSE), and CCoAOMT. The names of the enzyme families are indicated on the left of the trajectory. The enzyme (sub-)families present in a specific common ancestor have been plotted onto the respective nodes of the cladogram below the evolutionary scenario of the enzyme families involved in the phenylpropanoid pathway and lignin biosynthesis. White dots indicate absence/loss of a gene family one dot indicates the presence of one representative of the gene family and several dots indicate an expansion (two or more members of the gene family) in at least one species of the represented lineages in the cladogram. Colors are chosen to distinguish different enzyme families and subfamilies. Question marks label sequences of ambiguous affiliation.

Our data pinpoint that most of the enzymes have undergone massive lineage-specific expansions. A lineage-specific expansion is palpable even despite the fact that sampling of sequences across the Streptophyta is still strongly biased towards seed plants. These data offer a framework for pinpointing those candidate genes/enzymes that are bound to shed light on the evolution of key enzymatic steps—and novel ones. Such work is exemplified by studies on *Selaginella* or bryophyte model systems such as *Physcomitrium patens*. Characterizing enzymes that are even more divergent from what we know from angiosperms should yield surprising insights and novel routes in this bountiful pathway.

## MATERIAL AND METHODS

### Dataset of protein sequences and screening for homologs

We downloaded protein data from: (a) genomes of fifteen land plants: *Anthoceros agrestis* as well as *Anthoceros punctatus* (Li et al., 2020), *Amborella trichopoda* (*Amborella* Genome Project, 2013), *Arabidopsis thaliana* (Lamesch et al., 2010)*, Azolla filiculoides* (Li et al., 2018)*, Brachypodium distachyon* (The International Brachypodium Initiative, 2010)*, Capsella grandiflora* (Slotte et al., 2013)*, Gnetum montanum* (Wan et al., 2018)*, Marchantia polymorpha* (Bowman et al., 2017)*, Nicotiana tabacum* (Sierro et al., 2014)*, Oryza sativa* (Ouyang et al., 2007)*, Picea abies* (Nystedt et al., 2013)*, Physcomitrium patens* (Lang et al., 2018)*, Salvinia cucullata* (Li et al., 2018)*, Selaginella moellendorffii* (Banks et al., 2011), and *Theobroma cacao* (Argout et al., 2011); (b) the genomes of seven streptophyte algae: *Chlorokybus atmophyticus* (Wang et al., 2020), *Chara braunii* (Nishiyama et al., 2018), *Klebsormidium nitens* (Hori et al., 2014), *Mesotaenium endlicherianum* (Cheng et al., 2019), *Mesostigma viride* (Wang et al., 2020), *Penium margaritaceum* (Jiao et al., 2020), *Spirogloea muscicola* (Cheng et al., 2019); (c) the genomes of five chlorophytes: *Bathycoccus prasinos* (Moreau et al., 2012)*, Chlamydomonas reinhardtii* (Merchant et al., 2007), *Coccomyxa subellipsoidea* (Blanc et al., 2012), *Micromonas pusilla*, *Micromonas* sp. (Worden et al., 2009), *Ostreococcus lucimarinus* (Palenik et al., 2007), *Ulva mutabilis* (De Clerck et al., 2018), *Volvox carteri* (Prochnik et al., 2010). Additionally, we included sequences found in the transcriptomes of *Spirogyra pratensis* (de Vries et al., 2020), *Zygnema circumcarinatum* (de Vries et al., 2018), and *Coleochaete orbicularis* (Ju et al., 2015).

For each of the protein families we investigated here, the representative *Arabidopsis thaliana* protein was used as a query sequence for a BLASTp against this dataset. Initially, we considered all homologs recovered at a cutoff level of 10^-7^. However, due to the large size of the protein families (i.e. high number of well-supported homologs obtained), refinement of the datasets was carried out as described in the individual sections for these enzymes in the Results and Discussion section.

### Alignments, phylogenetic analysis, and primary sequence analysis

Using the homologs detected based on the above described BLASTp search for a given enzyme, we generated alignments using MAFFT v7.453 (Katoh and Standley, 2013) with a L-INS-I approach. Alignments were cropped, if necessary, to retain conserved domains that were alignable for all homologs; alignments are provided in Supplemental Datasets S1 to S11. We computed maximum likelihood phylogenies using IQ-TREE multicore version 1.5.5 (Nguyen et al., 2015), with 100 bootstrap replicates. To determine the best model, we used ModelFinder (Kalyaanamoorthy et al., 2017) and picked the best models based on the Bayesian Information Criterion. The best models were: LG+G4 (Le and Gascuel, 2008) for 4CL, CCR, CCoAOMT; LG+I+G4 for PAL, CAD, MAGL/CSE, COMT, and for the preliminary phylogeny of 4CL; LG+F+I+G4 for C4H, F5H, and C3H; WAG+F+G4 (Whelan and Goldman, 2001) for HCT.

Protein structure prediction was carried out using the sequences as input in the online Iterative Threading ASSEmbly Refinement (I-TASSER; Zhang, 2008; Yang et al., 2015). Functional residue analyses were based on published structural analyses (Rupasinghe et al., 2003; Ferrer et al., 2005, Youn et al., 2006; Hu et al., 2010; Pan et al., 2014) and alignments were viewed with SeaView v.4 (Gouy et al., 2009) and plotted with ETE3 (Huerta-Cepas et al., 2016).

### Protein domain predictions

Protein domains for all protein sequences for the enzyme families 4CL, CCR, C3H, CAD and CCoAOMT included in the phylogenies were predicted using InterProScan version 5.47-82.0 (Jones et al., 2014). The presence or absence of protein domains were mapped onto the phylogenies of the afore mentioned gene families as presence/absence heatmaps, which were visualized using iTOL v6 (Letunic and Bork, 2019).

## ACKNOWLEDGEMENTS

J.M.R.F.-J. is grateful for being supported by the Ph.D. program “Microbiology and Biochemistry” within the framework of the “Göttingen Graduate Center for Neurosciences, Biophysics, and Molecular Biosciences” (GGNB) at the University of Goettingen; A.D.A. is grateful for being supported through the International Max Planck Research School (IMPRS) for Genome Science. J.d.V. thanks the European Research Council for funding under the European Union’s Horizon 2020 research and innovation programme (Grant Agreement No. 852725; ERC-StG “TerreStriAL”). M.P., I.F., and J.d.V. are grateful for support through the German Research Foundation (DFG) within the framework of the Priority Programme “MAdLand – Molecular Adaptation to Land: Plant Evolution to Change” (SPP 2237; VR 132/4-1; PE 360/37-1; FE 446/14-1)

