## Supplementary material for "The evolution of the phenylpropanoid pathway entailed pronounced radiations and divergences of enzyme families": Figure S1

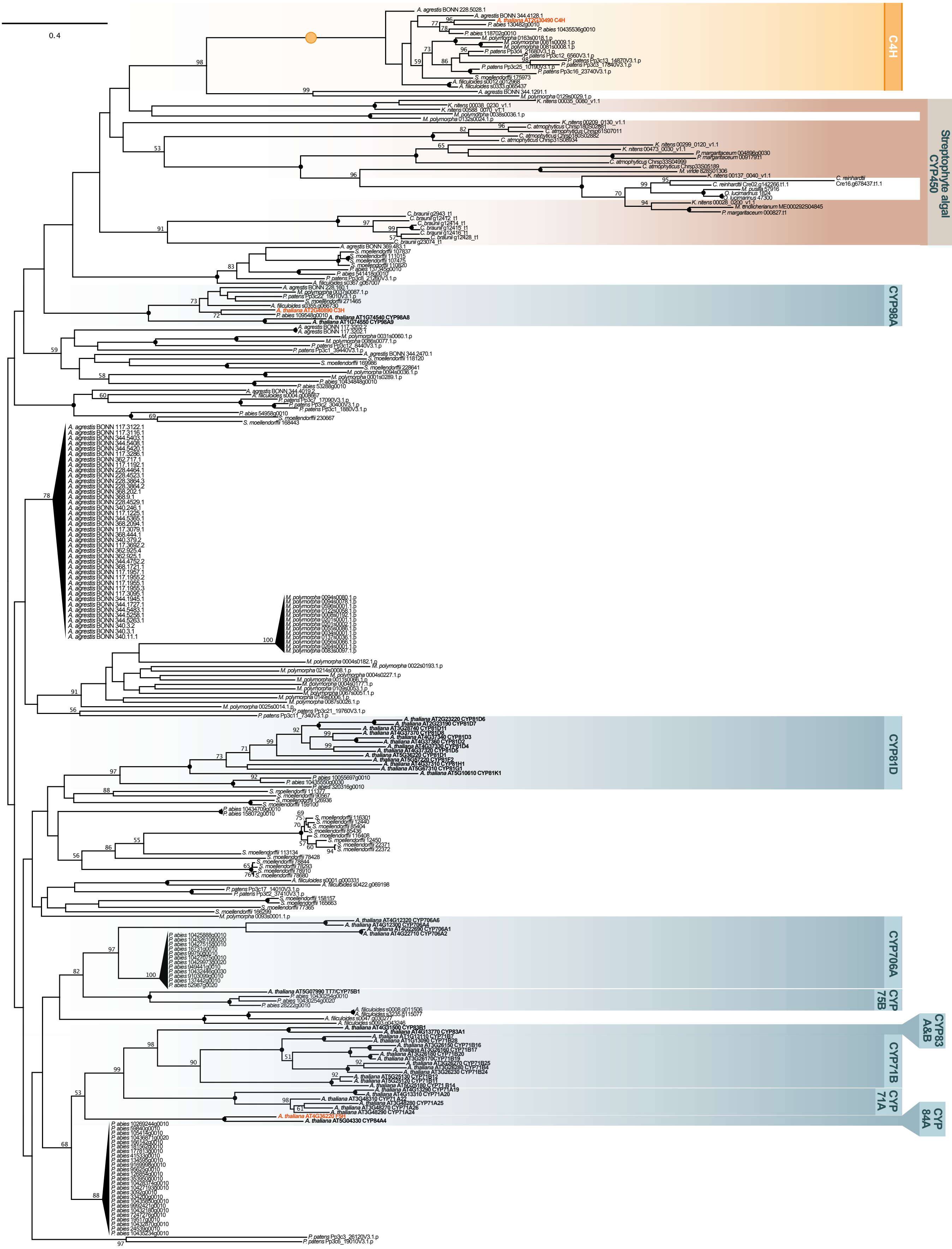


**Figure S1. Phylogeny of C4H homologs**. We sampled C4H homologs from seven land plant genomes that had a bit score of at least 200, as well as seven streptophyte algal and five chlorophyte algal genomes that had a bit score of at least 100. We aligned all C4H homologs and computed an unrooted maximum likelihood phylogeny using LG+F+I+G4 as model for protein evolution (chosen according to BIC). 100 bootstrap replicates were computed; only bootstrap values ≥50 are shown and bootstrap values of 100 are depicted by a filled dot. Colored font and dots correspond to the support recovered for the higher-order clades labeled on the right of the phylogenies.
