## Supplementary material for "The evolution of the phenylpropanoid pathway entailed pronounced radiations and divergences of enzyme families": Figure S2

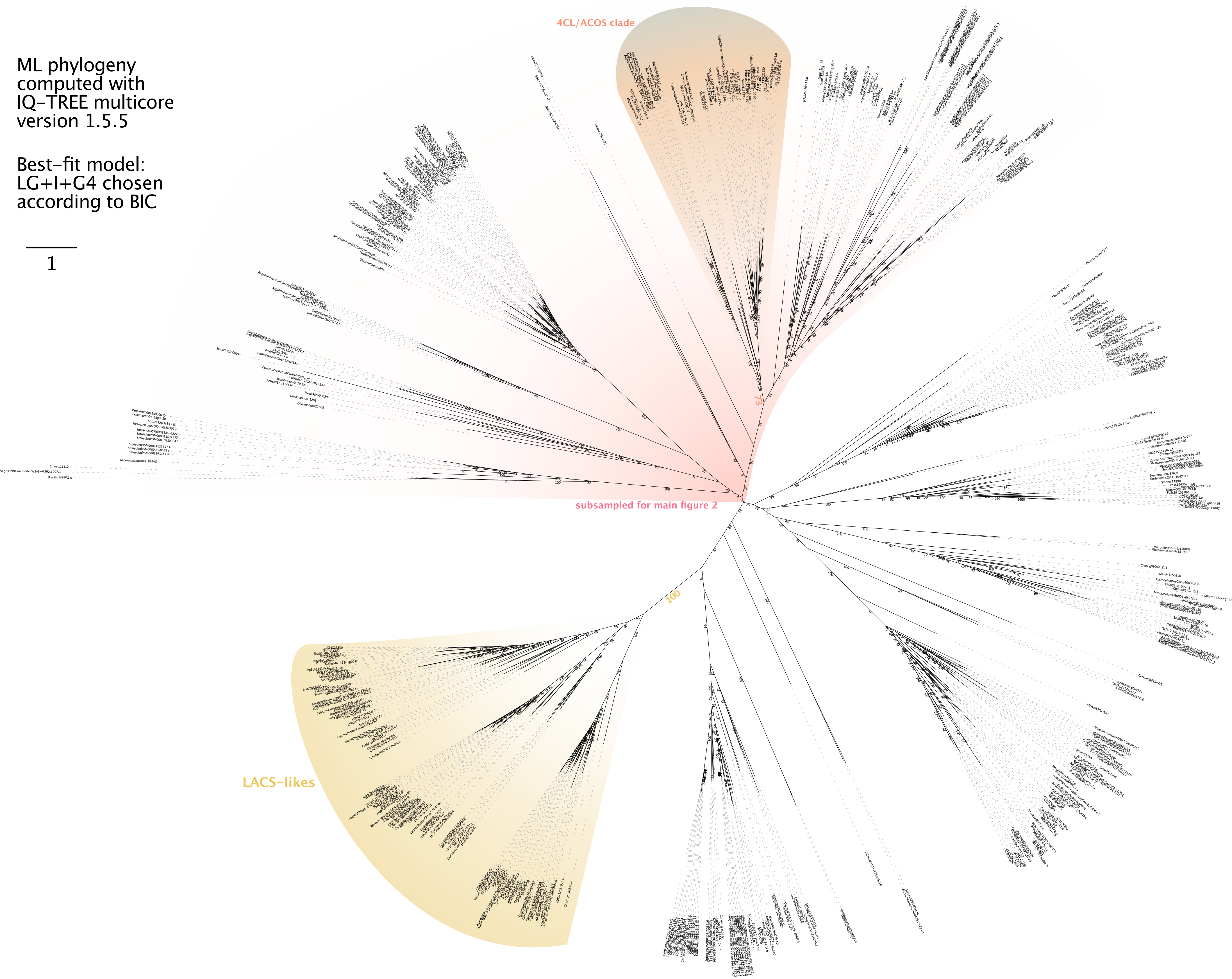


**Figure S2. Phylogeny of 4CLs / AAEs**. We sampled C4H homologs from nine land plant genomes, seven streptophyte algal and five chlorophyte algal genomes; additionally we added homologs found in transcriptomes of *Spirogyra pratensis* (de Vries et al., 2020), *Zygnema circumcarinatum* (de Vries et al., 2018), *Coleochaete scutata* (de Vries et al., 2018) and *Coleochaete orbicularis* (Ju et al., 2015). We aligned all 4CL/AAE homologs that had a length between 400 and 1150 aa and computed an unrooted maximum likelihood phylogeny using LG+I+G4 as model for protein evolution (chosen according to BIC). 100 bootstrap replicates were computed; only bootstrap values ≥50 are shown and bootstrap values of 100 are depicted by a filled dot. Colored font correspond to the support recovered for the higher-order clades; the pink gradient highlights which sequences/section of the tree was used for computing main Figure 3.
