## Supplementary material for "The evolution of the phenylpropanoid pathway entailed pronounced radiations and divergences of enzyme families": Figure S5

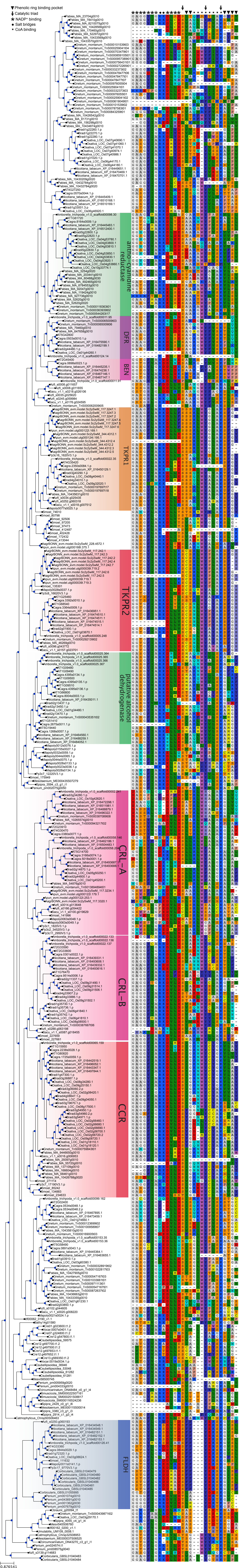


**Figure S5. Key residues of protein sequences in the phylogeny of CCR homologs**. The phylogenetic data on the left are identical to main Figure 4; relevant clade labelling was adapted from Figure 4. On the right (next to the leaves of the tree), we show key residues for the phenolic ring binding pocket, the catalytic triad, NADP$^+$ binding, salt bridges, and CoA binding of canonical CCR as reported by Pan et al. (2014).
