## Supplementary material for "The evolution of the phenylpropanoid pathway entailed pronounced radiations and divergences of enzyme families": Figure S7

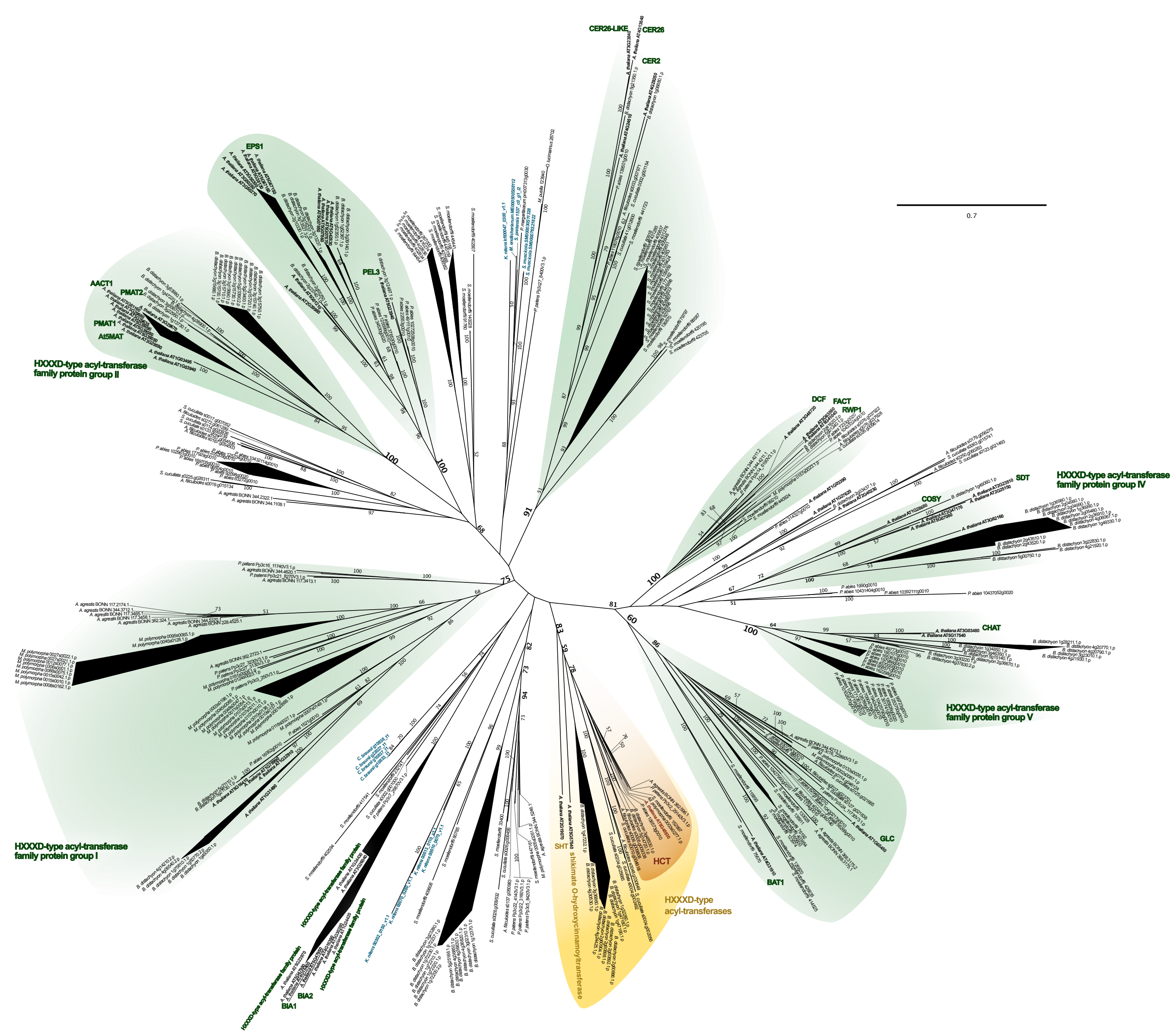


**Figure S7. Phylogeny of HCT homologs**. We sampled HCT homologs from fifteen land plant genomes, seven streptophyte algal and five chlorophyte algal genomes; additionally we added homologs found in the transcriptome of *Spirogyra pratensis* (de Vries et al., 2020). We aligned all HCT homologs that had a minimum lenght of 400 aa and computed an unrooted maximum likelihood phylogeny using WAG+F+G4 as model for protein evolution (chosen according to BIC). 100 bootstrap replicates were computed; only bootstrap values ≥50 are shown and bootstrap values of 100 are depicted by a filled dot. Colored font and dots correspond to the support recovered for the higher-order clades labeled on the right of the phylogenies.
