## Supplementary material for "The evolution of the phenylpropanoid pathway entailed pronounced radiations and divergences of enzyme families": Figure S8

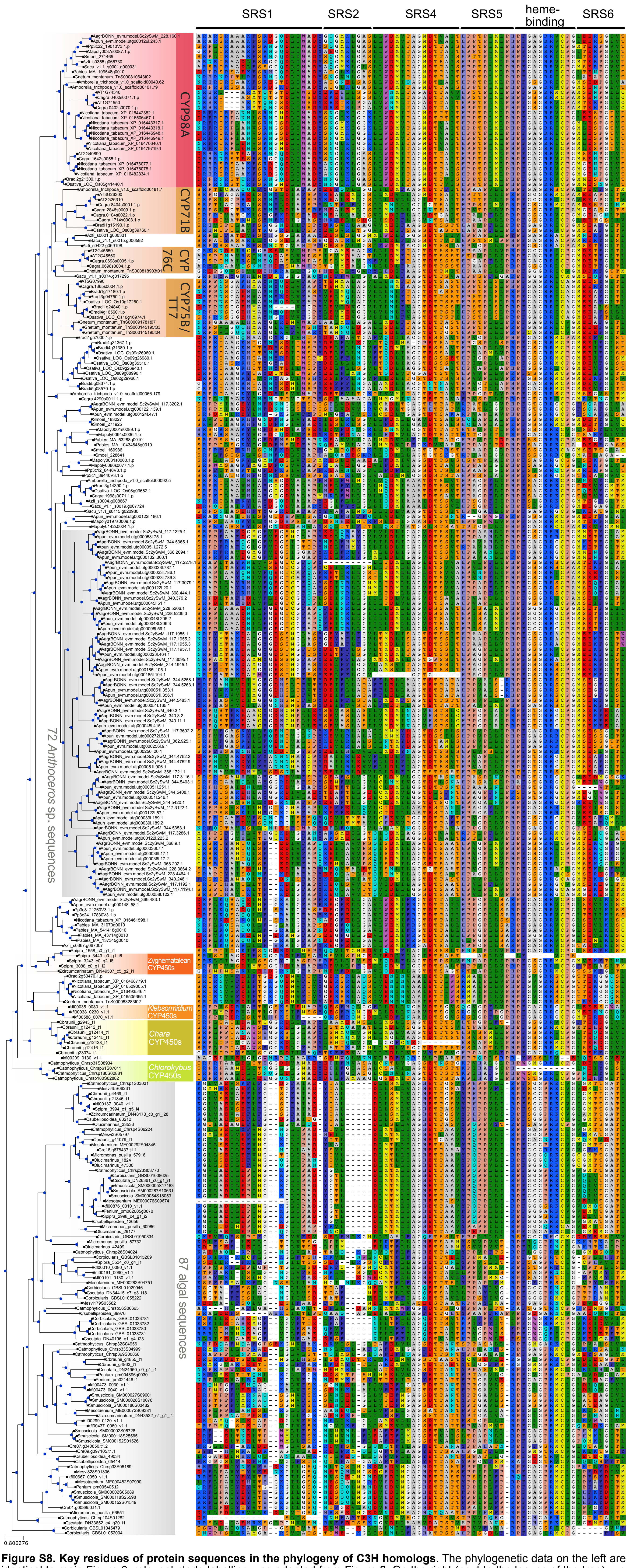


**Figure S8. Key residues of protein sequences in the phylogeny of C3H homologs**. The phylogenetic data on the left are identical to main Figure 6; relevant clade labelling was adapted from Figure 6. On the right (next to the leaves of the tree), we show key residues for the substrate recognition sites (SRS) as reported by Rupasinghe et al. (2003).
