## Supplementary material for "The evolution of the phenylpropanoid pathway entailed pronounced radiations and divergences of enzyme families": Figure S10

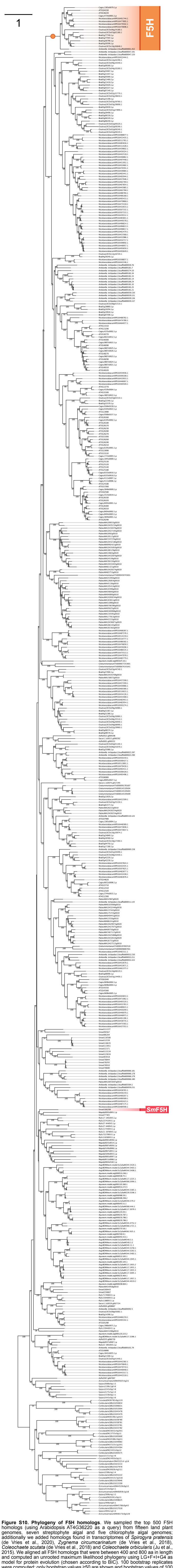


**Figure S10. Phylogeny of F5H homologs**. We sampled the top 500 F5H homologs (using Arabidopsis AT4G36220 as a query) from fifteen land plant genomes, seven streptophyte algal and five chlorophyte algal genomes; additionally we added homologs found in transcriptomes of *Spirogyra pratensis* (de Vries et al., 2020), *Zygnema circumcarinatum* (de Vries et al., 2018), *Coleochaete scutata* (de Vries et al., 2018) and *Coleochaete orbicularis* (Ju et al., 2015). We aligned all F5H homologs that were between 400 and 800 aa in length and computed an unrooted maximum likelihood phylogeny using LG+F+I+G4 as model for protein evolution (chosen according to BIC). 100 bootstrap replicates were computed; only bootstrap values ≥50 are shown and bootstrap values of 100 are depicted by a filled dot. Colored font and dots correspond to the support recovered for the higher-order clades labeled on the right of the phylogenies.
