## Supplementary material for "The evolution of the phenylpropanoid pathway entailed pronounced radiations and divergences of enzyme families": Figure S11

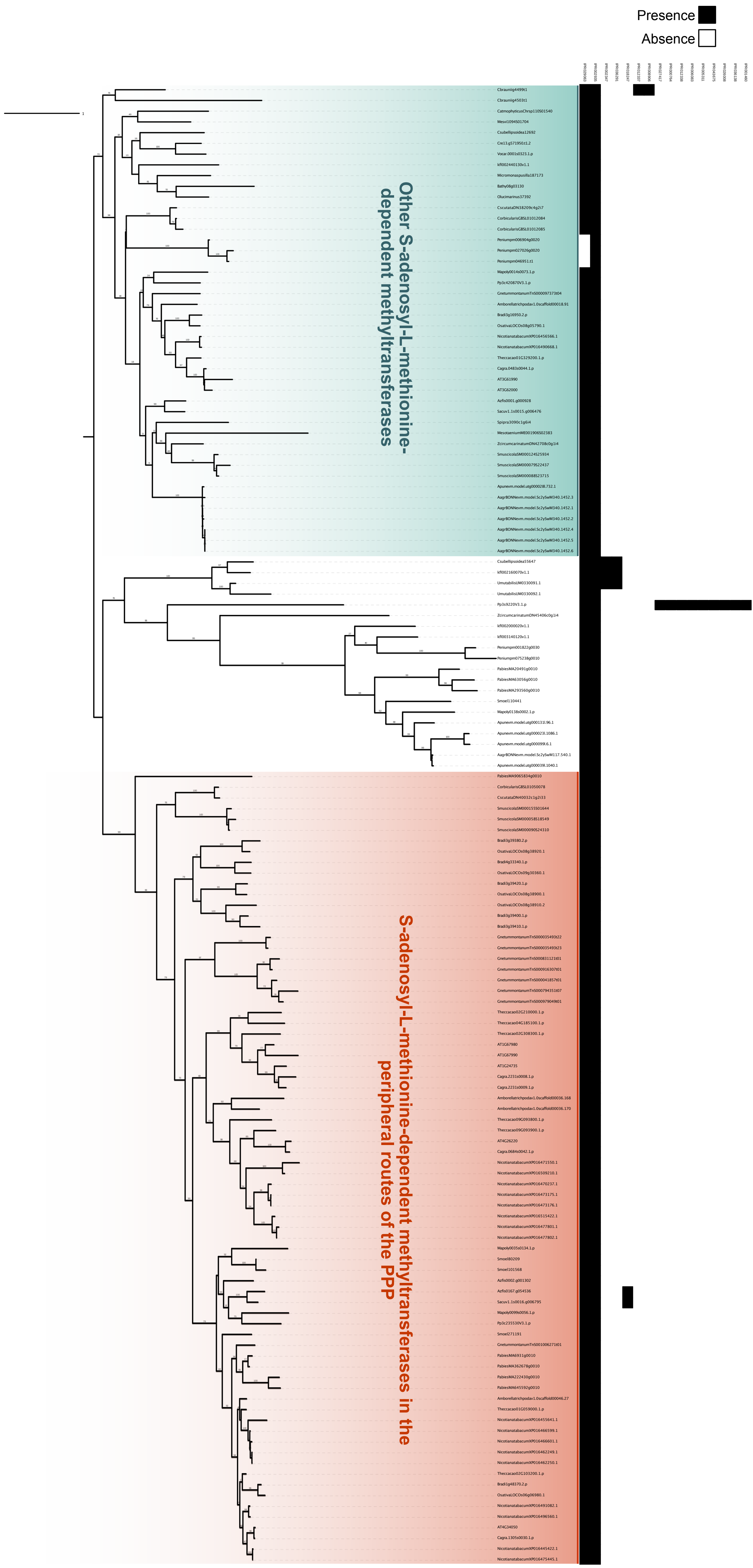


**Figure S11. Domains predicted for protein sequences in the phylogeny of CCoAOMT homologs**. The phylogenetic data on the left are identical to main Figure 9; relevant clade labelling was adapted from Figure 9. On the right (next to the leaves of the tree), black and white ticks indicate presence and absence of a given domain as predicted by InterProScan. Domain labels correspond to InterPro IDs.
