## Supplementary material for "The evolution of the phenylpropanoid pathway entailed pronounced radiations and divergences of enzyme families": Figure S12

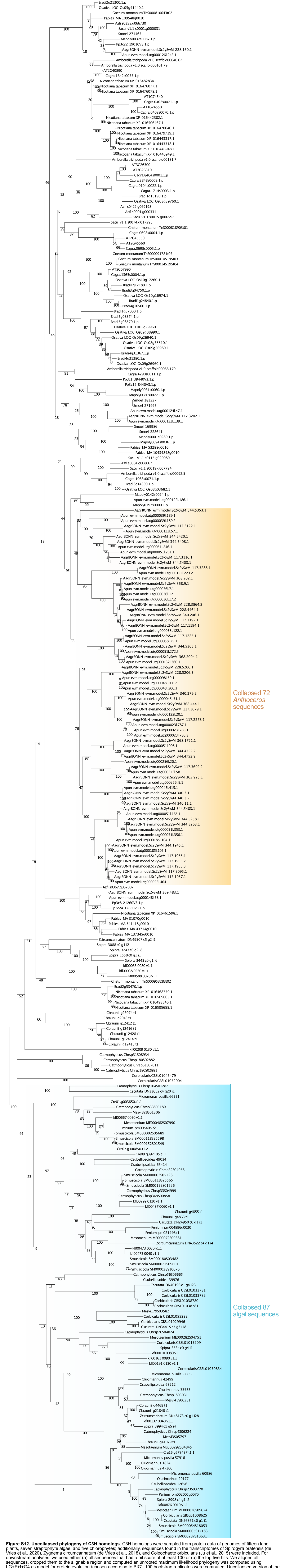


**Figure S12. Uncollapsed phylogeny of C3H homologs**. C3H homologs were sampled from protein data of genomes of fifteen land plants, seven streptophyte algae, and five chlorophytes; additionally, sequences found in the transcriptomes of Spirogyra pratensis (de Vries et al., 2020), Zygnema circumcarinatum (de Vries et al., 2018), and Coleochaete orbicularis (Ju et al., 2015) were included. For downstream analyses, we used either (a) all sequences that had a bit score of at least 100 or (b) the top five hits. We aligned all sequences, cropped them to the alignable region and computed an unrooted maximum likelihood phylogeny was computed using LG+F+I+G4 as model for protein evolution (chosen according to BIC). 100 bootstrap replicates were computed. Uncollapsed version of the C3H phylogeny shown in Figure 6; colored regions highlight the clades that were collapsed in Figure 6.
